# Mapping gut bacteria into functional niches reveals the ecological structure of human gut microbiomes

**DOI:** 10.1101/2023.07.04.547750

**Authors:** Laura Anthamatten, Philipp Rogalla von Bieberstein, Clémentine Thabuis, Carmen Menzi, Markus Reichlin, Marco Meola, Bertrand Rodriguez, Otto X. Cordero, Christophe Lacroix, Tomas de Wouters, Gabriel E. Leventhal

## Abstract

Microbiomes are an essential contributor to the metabolic activity in the human gastrointestinal tract. The fermentation of otherwise indigestible nutritional components like dietary fibers relies on a complex interplay of metabolic pathways that are distributed across the individual bacteria. Yet, which of the bacteria are responsible for which parts of the distributed metabolism and how they should be grouped together is insufficiently understood. Here, we present the NicheMap™, an approach to map the different bacterial taxa that make up the gut microbiome onto the different functional niches of microbial carbohydrate fermentation. Our approach uses *in vitro* measurements of bacterial growth and metabolic activity to identify which bacterial taxa are responsible for which metabolic function in the relevant complex context of whole human fecal microbiomes. We identified ‘characteristic taxa’ selected for by a panel growth substrates representative of dietary components that are resistant to digestion by host enzymes. These characteristic taxa offer predictions of which bacteria are stimulated by the various components of human diet. We validated these predictions using microbiome data from a human nutritional supplementation study. We suggest a template of how bacterial taxonomic diversity is organized along the trophic cascade of intestinal carbohydrate fermentation. We anticipate that our results and our approach will provide a key contribution towards building a structure-function map for gut microbiomes. Having such a map on hand is an important step in moving the microbiome from a descriptive science to an interventional one.

---

The human gut microbiome contributes to physiology and health through its metabolic activity—breaking down otherwise indigestible components of our diet into metabolites that affect the host—for example, shortchain fatty acids^1, 2^. The breakdown of these complex carbohydrates is typically not performed by any individual microbe alone, but rather requires metabolic pathways that are distributed across the many different microbial constituents of the microbiome^3^. If we want to therapeutically modify the microbiome, we thus need to know (i) how species abundances differ between health and disease, and (ii) what aspect of the distributed function each of the species is responsible for. While measuring microbiome composition has become widely accessible through genomic sequencing, directly measuring the contribution of individual microbes to microbiome function in its relevant context is still an unsolved challenge. One solution to this challenge is to create a map of taxato-functions that can translate microbiome composition to microbiome function. Yet, no such mapping exists, and this gap is one of the main obstacles for moving microbiome science from a descriptive discipline to an interventional one.

Exhaustively probing all species or strains in the gut microbiome is prohibitive, both because of the large genotypic diversity of gut microbes, but also because microbe-microbe interactions and microbe-host interactions modulate microbial metabolic activity. Ecological theory is often invoked to provide a conceptual framework for complexity reduction^4, 5^. One such concept is that of functional groups or guilds^6^, where functionally equivalent taxa in terms of the resources they consume are grouped together to simplify the system and describe it in functional instead of taxonomic terms^7, 8^. Through this lens, microbiomes assemble with one taxon per functional guild because of strong resource competition between members of the same guild^9^.

Although the idea of guilds is conceptually appealing, its quantitative implementation is not straightforward. First, ‘function’ can be defined in multiple and nonmutually exclusive ways (e.g. the presence of a metabolic pathway, the ability to produce a secondary metabolite, etc.). Second, most taxa have the capability to perform a broad range of functions, but there is important context-dependence in terms of which biotic and abiotic environments the microbes express these functions in^10^. Thus, even once functions are defined, measuring them in their relevant contexts is non-trivial. Because of these difficulties, most work to date defines function at a genomic level that can then be profiled in high throughput^11, 12^. But large uncertainties in genomic functional prediction—in particular for non-model bacteria—dampen the translational utility of these approaches.

Here, we present an approach to circumvent these difficulties and construct a bacteria-to-function mapping based on direct *in vitro* phenotypic measurements of complex human fecal microbiomes that we call the NicheMap™. We look at the microbiome from a metabolic perspective and essentially view the gut microbiome as an anaerobic digester of dietary fibers^13^. The advantage is that the key microbiome functions derive directly from the well-defined biochemistry of microbial fermentation. From this point of view, a microbiome metabolically encodes a cascade of hydrolytic and fermentative pathways that convert complex carbohydrates down to simple metabolic end-products—mainly organic acids—which then further interact with the host^14–16^. Specifically, we performed growth competition experiments within whole human fecal microbiomes in strict anaerobic batch fermentations using diluted fecal samples from eight human donors as inocula. We replicated these experiments by supplementing each enrichment with one of nine different growth substrates that are representative of the components of human diet that are resistant to digestion by host enzymes. By measuring (a) the changes in microbial composition together with (b) the produced metabolites after 48 h of cultivation, we were able to place ‘characteristic taxa’ into metabolic niches that map back to the trophic fermentation cascade. We then asked which of these niches are structured as functional guilds and which are taxonomically conserved with variable functional output because of context-dependence. Finally, we validated our predictions with microbiome data from a nutritional intervention study. Our NicheMap™ platform thus puts forth an approach to functionally characterize microbiome bacteria and establish a taxon-function-map that takes into account the relevant microbiome context.

## Results

### The total metabolic output is more strongly determined by carbon source than microbiome composition

Our premise is that competition for the different extrinsic and intrinsic resources ultimately determines which bacteria thrive and are metabolically active in the gut. Extrinsic resources comprise those parts of the diet that are not digested or absorbed prior to arrival in the relevant portion of the human gastrointestinal tract (typically the colon), as well as different host and microbial glycans, peptides, and other nutrients. Focusing on dietary fibers, these are hydrolyzed into simpler building blocks and converted via fermentative pathways into different fermentation products. The most frequent fermentation intermediate products are formate, lactate, and succinate, and typical fermentation end products are acetate, butyrate, and propionate (Figure 1a). We experimentally simulated the outcomes of competition for extrinsic and intrinsic resources using diluted fecal samples from eight different human fecal donors in a defined base medium supplemented with a panel of substrates that are representative of the extrinsic dietary components subject to colonic fermentation: fibers (arabinogalactan, AG; pectin, PE; pea fiber, PF; resistant dextrin, RD), simpler polysaccharides (xylan, XY; soluble starch, SS), and glycosylated proteins (mucin, MU; yeast extract, YE). We standardized the media to contain 3 g*/*L of extrinsic carbon, corresponding to roughly 100 mM supplemented carbon atoms for each of the substrates (see Supplementary Table S1). To cover a broad range of possible bacterial responses, we chose fecal donors with microbiome compositions that span the entire enterotype diversity with respect to *Bacteroides*, *Prevotella*, and *Ruminococcus* (Supplementary Figure S1).

**Figure 1:**
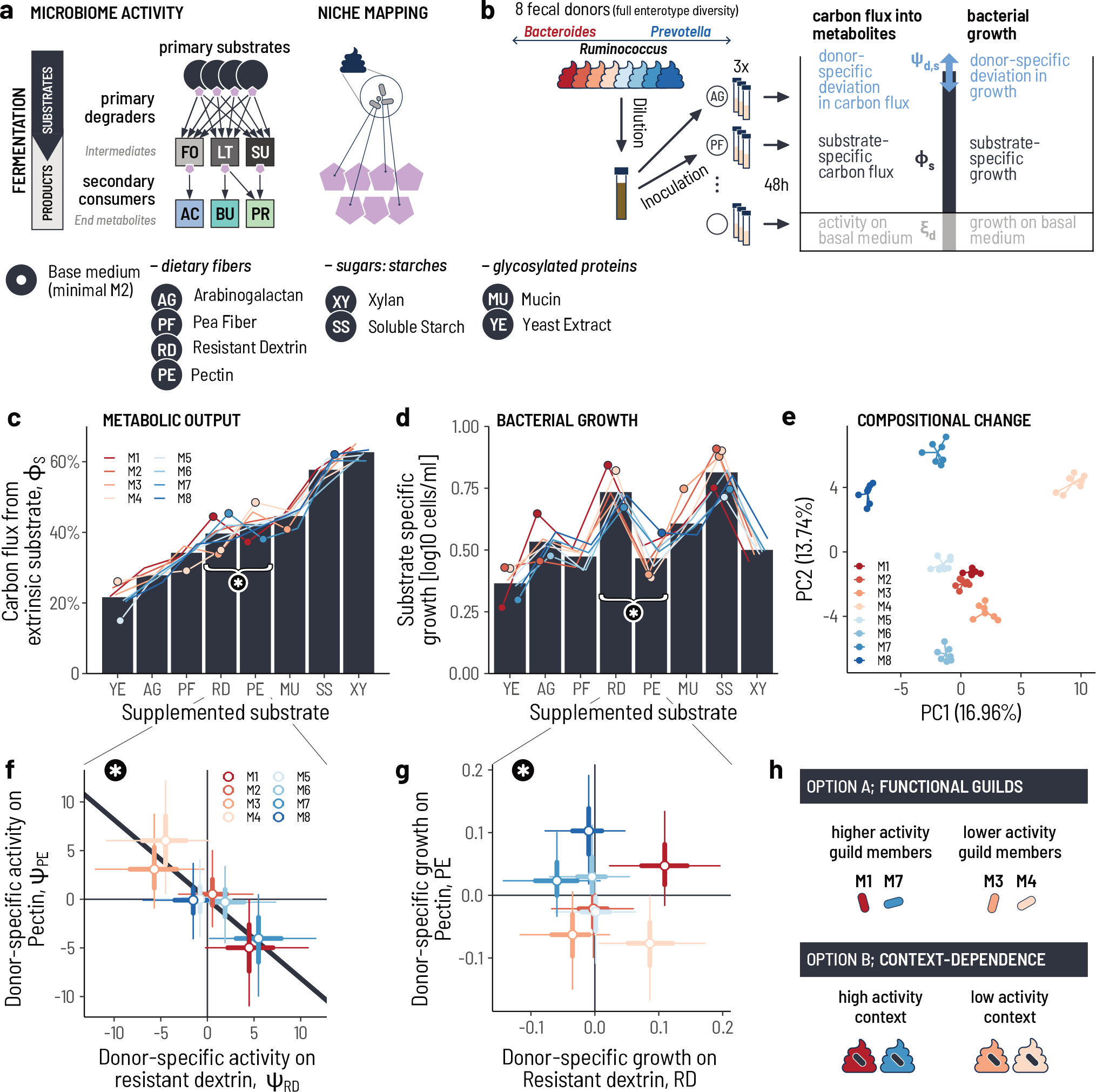
Microbiome activity is more strongly determined by growth substrate than microbiome identity. **a.** The goal is to map gut bacteria into the functional niches of the gut. We use carbon fermentation as a biochemical scaffold, where each primary resource (e.g. dietary fibers) or secondary resource (e.g. fermentation intermediates) represents a niche. **b.** Whole fecal microbiota that span the entire enterotype spectrum are enriched in a basal medium that is supplemented with a panel of complex carbon sources. Metabolites are measured after 48 h of activity, and the composition is determined with 16S amplicon sequencing. We statistically partition the readouts into three components: the activity from the basal medium, the activity specific to the carbon source, the donor-substrate specific deviation. **c.** The carbon flux into the measured metabolites varies by supplemented substrate. The bars show the estimated mean across all donors, and the colored lines show the variation of individual donors. Circles indicate donor/substrate pairs that are significantly different from the respective mean (posterior probability < 0.1). **d.** The substrate-specific bacterial growth [cells/ml] between substrates. The bars show the posterior mean increase in biomass across all donors, and the colored lines show the individual donors. Circles indicate donors for which the biomass increase is significantly different from the mean (posterior probability > 0.1). The ordering of the substrates is the same as in panel c. **e.** The eight microbiota retain a strong compositional signature of the fecal sample after enrichment. Each point shows the top two principal components (PC) of the bacterial composition at the end of an enrichment from a donor fecal inoculum in an individual growth substrate. **f.** Donor microbiota M1 and M8 are particularly efficient at metabolizing resistant dextrin (RD) and are comparatively poor at metabolizing pectin (PE), and vice-versa for M3 and M4. The circles show the posterior means, the thick lines the posterior 50% and the thin lines the 95% highest probability density intervals. **g.** The observed metabolic trade-off between RD and PE is not recapitulated for total bacterial growth. **h.** We postulate that the observed commonalities and differences in total metabolic activity and bacterial growth can be mapped back to either (A) the occurrence of functional guilds, wherein each fecal microbiome might have its own specific guild member, or (B) that the niches are filled by the same bacteria across fecal donors, but that the remaining biotic context modulates the activity and growth.

With this setup at hand, we first asked whether the tested microbiomes differed in terms of the realized carbon flux, that is, their overall capacity to convert the primary substrates into fermentation products. We estimated the carbon flux from each of the extrinsic carbon sources using a Bayesian approach (see Methods). Briefly, we assumed a total of 100 mM supplemented carbon for all substrates and computed the proportion of the carbon that ended up in the measured metabolites formate, lactate, succinate, acetate, butyrate, propionate, and ethanol. We then partitioned the carbon flux into three components (Figure 1b): (i) the carbon flux from the basal medium for a donor *d*, *ξ_d_*; (ii) the carbon flux from the specific extrinsic substrate *s*, *ϕ_s_*; and (iii) the deviation from the mean carbon flux on substrate *s* for each donor *d*, *ψ_d,s_*. High values of *ϕ_s_* imply extrinsic substrates that are readily metabolized by the fecal microbiota, while low values suggest substrates that are more recalcitrant. Conversely, a high or low value of *ψ_d,s_* identifies a donor microbiota *d* that is particularly efficient or poor at metabolizing the extrinsic substrate *s*, respectively, compared to the average donor microbiota.

The substrates differed strongly in terms of the mean carbon flux, *ϕ_s_*(Figure 1c) that was achieved after 48 h. The highest carbon flux was from the structurally simple polysaccharides, soluble starch (simple glucose polymers; *ϕ*_SS_ = 55.5 %) and xylan (simple xylose polymers; *ϕ*_XY_ = 60.6 %). The carbon flux from the dietary fibers decreased with increasing structural complexity: pectin (complex soluble mix of rhamnose and galactose) and resistant dextrin (glucose polymers with resistant osidic linkages) were the highest among fibers (*ϕ*_PE_ = 42.4 %, *ϕ*_RD_ = 39.5 %, followed by pea fiber (soluble and insoluble fibers including pectin, cellulose, and xyloglucans; *ϕ*_PF_ = 33.7 %) and arabinogalactan (mix of arabinose and galactose; *ϕ*_AG_ = 27.3 %). This hierarchy of the substrates follows the expectation that structural complexity and solubility influences fermentability^17^.

The identity of the microbiota had a much smaller effect on carbon flux than the type of supplemented substrate. Individual donor microbiota deviated from the mean carbon flux between *−*6.6 % (M5 on YE) to 6.1 % (M4 on PE), implying that the donor microbiota all had similar capacities to metabolize the different extrinsic substrates. This was also true in terms of their capacity to metabolize the basal medium (Supplementary Figure S2). Hence, despite strong differences in composition across the tested microbiota, the overall functional output was largely conserved.

We asked whether the large differences in realized carbon flux between substrates (up to 3×) was the result of differences in bioavailability between the substrates, or alternatively the result of a trade-off between catabolism and anabolism, where lower carbon fluxes into metabolites implied a higher carbon flux into growth. We thus estimated the amount of bacterial growth across the different supplemented substrates as cells/ml based on the extracted DNA concentration calibrated to total cell counts with qPCR (see Methods). We then used the same partitioning approach into mean substrate effects and donor-specific deviations as for carbon flux (Figure 1b). Lower bioavailability implies that those substrates with lower carbon flux also had lower biomass yield. In contrast, a shift from metabolite production to biomass production implies that substrates with high carbon flux into metabolites would have lower biomass production, and *vice versa*.

The different primary substrates differed substantially in terms of how much bacterial growth they supported (Figure 1d), but the ranking did not correlate with that for carbon flux into metabolites (Spearman’s *r* = 0.476, *p* = 0.243). While some substrates with high carbon flux into metabolites also had a high biomass yield (e.g. SS, 6.5-fold increase in biomass over the basal medium), others did not (e.g. XY, 3.2-fold). Conversely, some substrates with intermediate carbon flux had particularly high biomass yields (RD, 5.4-fold) while others had lower yields (PE, 2.9-fold). Overall, there was no consistent mapping of carbon flux to growth, but rather each substrate was characterized by its own unique allocation into metabolite production and growth.

The unique signature of carbon flux and growth in each substrate suggested that different biochemical processes (enzymes, pathways, etc.) might be at play—and potentially also different taxa. We hypothesized that if the same taxa were responsible for metabolizing a substrate across donors, then the microbiome compositions at the end of each of the enrichments should converge and cluster by substrate more strongly than by donor. However, the different fecal donor microbiota strongly retained their initial differences in composition throughout the enrichments (Figure 1e). Thus, either the donor-specific compositional signal is much stronger in magnitude than the changes of the specific taxa that are responsible for metabolising a substrate, or otherwise each donor has its own specific taxon—or group of taxa—for each functional role.

We observed some notable differences between donor microbiota in specific cases (circles in Figure 1c,d; Supplementary Figure S3). Most of these significant donor-specific deviations pertained to either RD or PE, with an apparent trade-off between the two (asterisk in Figure 1c). Microbiota with a high carbon flux in RD had a low flux in PE (e.g. M7 and M1), and conversely, microbiota with a high flux in PE had a low flux in RD (Spearman’s *r* = *−*0.91, *p* = 0.001; Figure 1f, Supplementary Figure S4). This suggests that there are indeed properties of microbiome composition that make it more proficient in one function, but less so in another. The trade-off between RD and PE, however, was not recapitulated for bacterial growth (Figure 1g; Spearman’s *r* = *−*0.09, *p* = 0.84), and more generally carbon flux did not trade-off with bacterial growth, expect for PE (Spearman’s *r* = *−*0.81, *p* = 0.0218).

The structural complexity of the substrate was thus the main determinant of carbon flux into metabolites. Nevertheless, some yet uncharacterized donor-specific effects led to differences in bacterial growth and metabolic activity. To help explain these observations, we turned to two principles from ecological theory (Figure 1h). First, different bacteria could be responsible for the same function and therefore might be grouped into functional guilds. Small differences in how these bacteria express the function could then result in subtle differences in carbon flux and growth. Second, the same taxa could be responsible for a certain function, but because of differences in the background composition—i.e. the microbiome context—the breakdown products are metabolized in a slightly different way, again resulting in subtle differences in terms of carbon flux and growth. To investigate which of the two scenarios was more consistent with our data, we turned to the compositional changes that occurred during the enrichments.

### Each growth substrate has characteristic taxa that are mostly conserved across microbiota

We first verified that the basal culture conditions themselves did not bias bacterial growth unevenly across donor fecal samples (Supplementary Text). Next, we identified putative ‘characteristic taxa’ that are drivers of the differential activity signals across substrates by determining two quantities for each taxon: (i) the degree of selection in substrate *s*, *A_s_*, and (ii) the absolute growth, *B_s_*. We estimated *A_s_* as the log_2_-fold increase in absolute abundance in a specific substrate relative to the basal medium, and *B_s_* as the difference in absolute abundance between enrichments in substrate *s* and the basal medium without supplemented substrate (Figure 2a). To reduce the compositional complexity, we grouped all the 1579 amplicon sequence variants (ASVs) into 231 phylogenetically coherent genera (GTDB taxonomy; see Methods).

**Figure 2:**
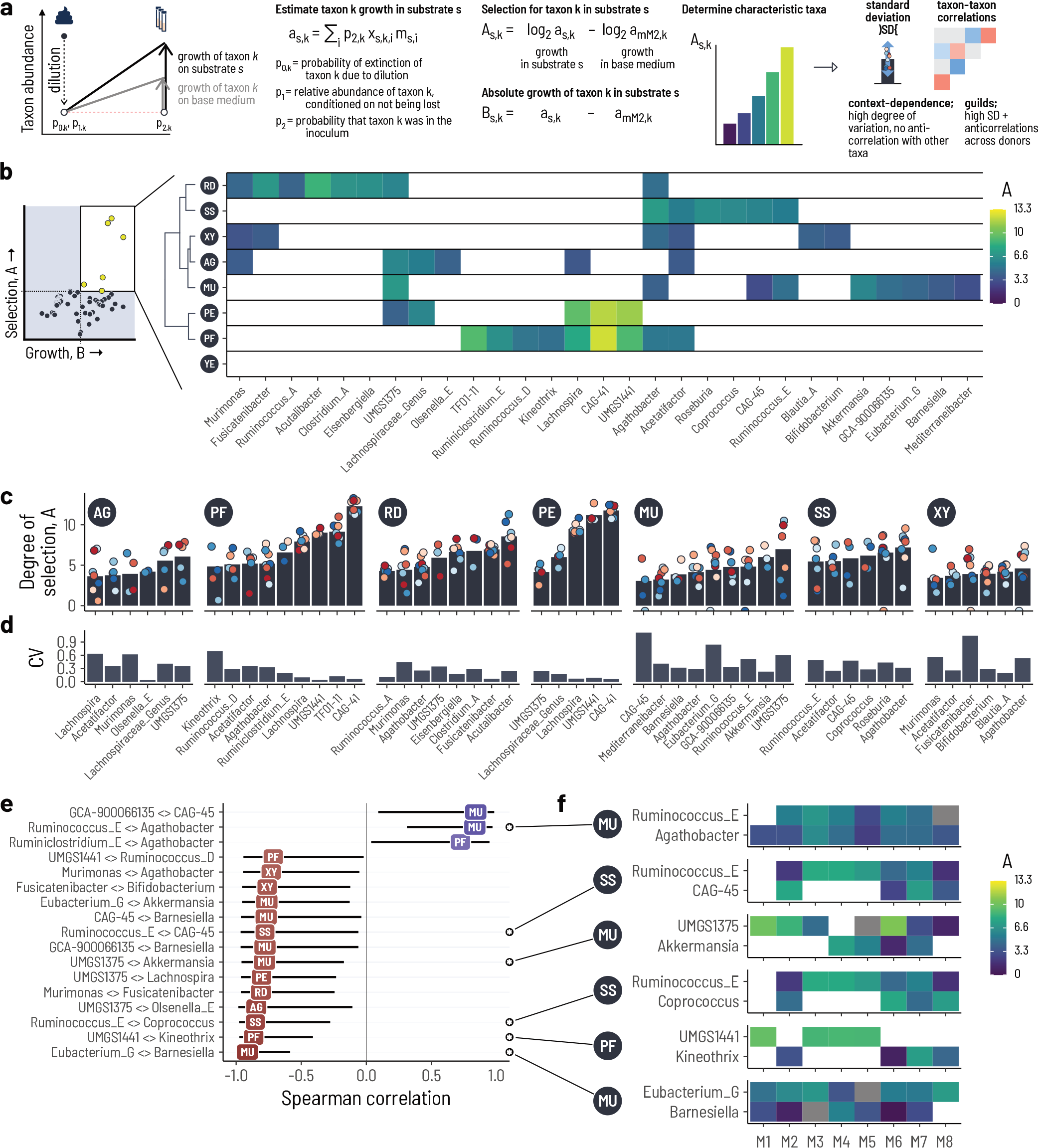
The primary substrates each select for specific characteristic taxa. **a.** Overview of the approach. We compute the selection for a specific taxon *k*, *A_s,k_*, and the absolute growth, *B_s,k_* in a substrate *s*. The stochastic effects from the initial dilution are accounted for probabilistically by computing *p*_0_, *p*_1_, and *p*_2_. *x_s,k,i_* is the relative abundance of taxon *k* in replicate *i* of substrate *s*. *m_s,i_* is the estimated total bacteria of replicate *i* in substrate *s*. Based on these values, we identify characteristic taxa that are specifically selected for on a substrate. We then use the variation in degree of selection *A* across donor microbiota and correlations between taxa to interpret how much context dependence and/or guild structure plays a role. **b.** The characteristic genera for a substrate are above a threshold for unspecific growth and corresponding selection. Most characteristic genera are specific to one (or two) substrates, though some are selected for across almost all substrates (marked with #). **c.** The bars show the mean selection across all donors and the circles show the respective values in a single donor. **d.** The bars show the coefficient of variation (CV) of *A* across donors. e. Only few correlations between the characteristic taxa within a substrate are significant. f. Distribution of *A* values for four examples of the taxon-taxon correlations.

The mean absolute growth of genera across substrates had a bimodal distribution with a separation between the two peaks at around *B_T_* = 5 *×* 10^6^ cells*/*mL (Supplementary Figure S5). We interpreted the lower peak as unspecific growth and defined *B_T_* as a cut-off for specific growth. For each substrate, we then determined the genera that were more strongly selected for than those below the threshold *B_T_* and were expected to be in the inoculum of at least two donors (*n* = 86; Figure 2b, Supplementary Figure S6). For YE, no genera were above the cutoff. Because the basal medium already contains yeast extract as a source of essential amino acids, cofactors, etc., additionally supplementing additional yeast extract likely did not specifically select for additional taxa. The *A* values of the characteristic taxa did not correlate with the abundance of the respective genera in the inoculum samples (Spearman’s *r* = *−*0.0468, *p* = 0.422) and thus we concluded that these were not *a priori* biased by differences in starting abundances. Overall, this procedure identified 29 out of 86 genera as characteristic for at least one substrate.

Each substrate had its own distinct set of characteristic genera. Most of the characteristic genera were only associated with one or two substrates (24/29; Figure 2b). The strongest selection occurred on PE and PF and these substrates overlapped in their characteristic genera, with the genus cluster CAG-41 (*Clostridia* class) and UMGS1441 (*Lachnospirales* order) among the top 3 in both substrates (Figure 2c). This overlap in characteristic taxa likely reflects the similarity between the two substrates, as PF contains pectins and the known pectin-utilizing genus *Lachnospira* (e.g. *L. eligens*, *L. pectinoschiza*) was also strongly selected for in both PE and PF^18^. For RD, the characteristic genera were particularly unique, including *Acutalibacter*, *Clostridium_A*, and *Eisenbergiella*.

The characteristic taxa we identified included those that are expected from the literature. MU selected for *Akkermansia* and *Barnesiella*, both canonical mucin degraders^19^, SS selected for the starch-utilizers *Agathobacter*, *Roseburia*, *Coprococcus*, and *Ruminococcus_E* (i.e. *R. bromii*)^20^, and XY selected for *Bifidobacterium* amongst others^21^.

Having identified the characteristic taxa for each of the substrates, we asked whether they exhibited signatures of functional guilds and/or context dependence. On the one hand, if different characteristic taxa were to make up a functional guild, then at least two of them should have both considerable variation and a negative correlation in their *A* values across donors. On the other hand, if the variation in metabolic output is driven by context dependence, then there should be few characteristic taxa per substrate and these should not have any anti-correlation with the other taxa (Figure 2a).

The characteristic genera with the highest *A* values were the most consistent across fecal donors. The coefficient of variation (CV) was generally lower than 1 (Figure 2d), implying that the amount of variation was lower than the magnitude of selection. The CV also generally decreased with increasing *A* (Spearman’s *r* = *−*0.583, *p* = 1.08 *×* 10*^−^*^5^; Supplementary Figure S7). This was particularly evident for PE, PF, and RD, underlining the specificity of fiber degradation. Nevertheless, some of the specific characteristic genera had high CVs compared to the others (e.g. *Acutalibacter* on RD, or UMGS1375 and *Akkermansia* on MU). To test whether this variation implied trade-offs with other taxa, we computed the pairwise correlations between all characteristic genera within a substrate.

Only few pairs of characteristic strains within a substrate correlated significantly in terms of *A*. Of the tested 155 pairs, 17 yielded correlations with *p <* 0.05, and 14 of those were negative (Figure 2e, Supplementary Figure S8). Most of these pertained to MU (7/17). Importantly, negative correlations mostly involved the lower ranked characteristic genera (e.g. *Eubacterium_G* and *Barnesiella* in MU, UMGS1441 and *Kineothrix* in PF, *Ruminococcus_E* and *Coprococcus* on SS; Figure 2f). A notable exception is *Akkermansia* and UMGS1375 in MU, which show signs of a stronger trade-off (Figure 2f). Mostly, however, the dominating characteristic taxa were the same across fecal donors, and putative guilds involving sub-dominant characteristic taxa.

### The dominant characteristic taxa are predictive of the metabolic output of the microbiota

With the characteristic taxa at hand, we next asked whether the metabolic activities of these bacteria can explain the overall community metabolic output on the different substrates (Figure 3a). To this end we selected two pure isolates that matched the characteristic taxa of RD and PE. For RD, we isolated an *Acutalibacter* sp. (strain PB-SMJER) after enrichment of a fecal sample from donor M7 on RD. For PE, we selected a *Lachnospira eligens* (strain PB-SJATG) that had previously been isolated from a fecal sample that was enriched in a medium high in pectins. We then measured the concentration of metabolites in terms of moles carbon produced by these two ‘characteristic strains’ on PE and RD, respectively.

**Figure 3:**
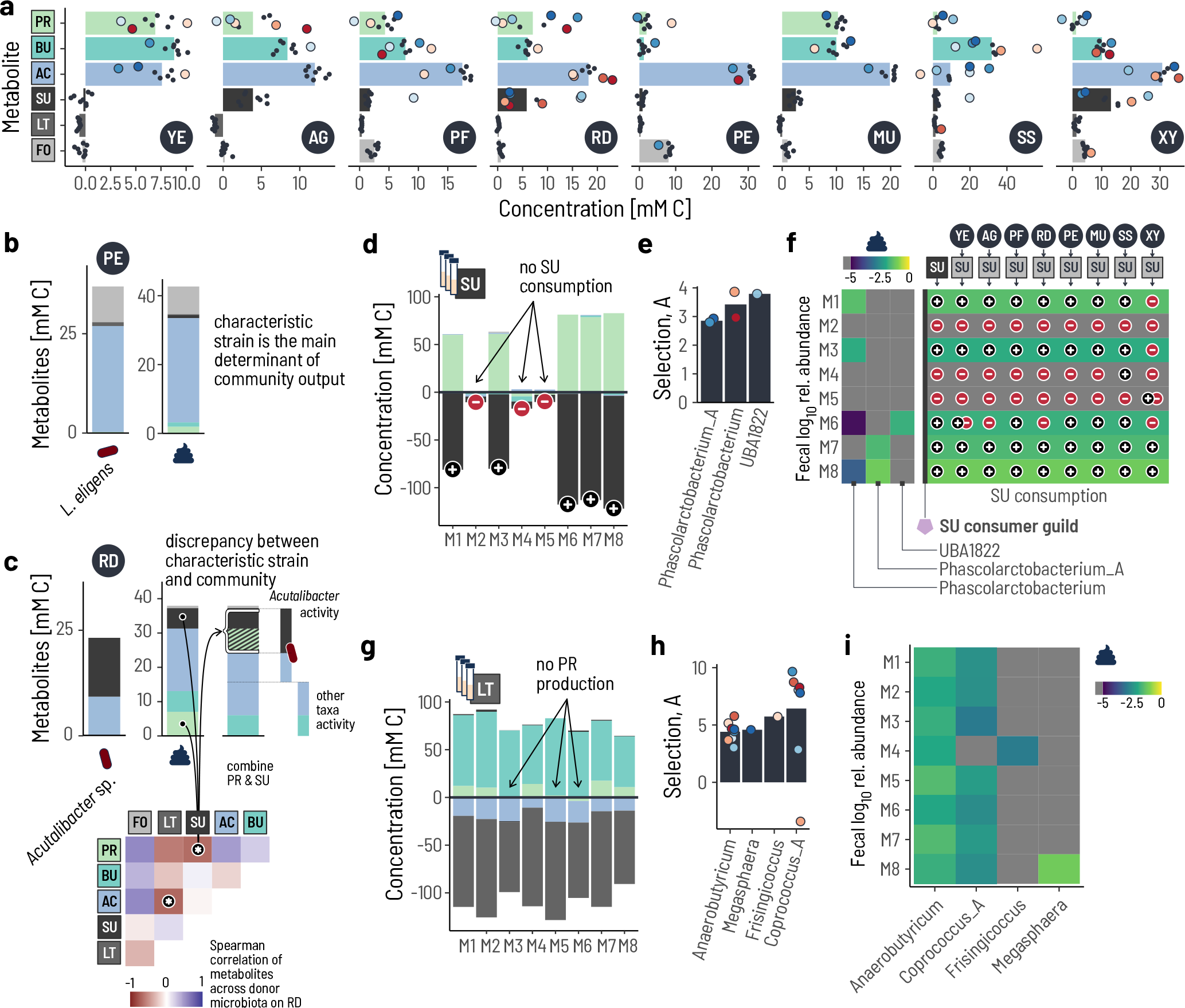
The metabolic activity of the characteristic taxa predicts that of the whole community. **a.** The metabolic output differs by added substrate. The bars show the mean concentration of the respective metabolite across all donors. The large colored circles indicate donor-specific deviations that are significantly different from zero (posterior probability > 0.1). The small black dots show the other donors. Concentrations are shown in mM carbon. FO: formate, LT: lactate, SU: succinate, AC: acetate, BU: butyrate, PR: propionate. The metabolite colors apply to the whole figure. **b.** The metabolic output of the characteristic taxon for pectin (PE), *Lachnospira eligens*, matches the metabolic output of the whole community. *L. eligens* was in monoculture on the PE-supplemented medium in the same conditions as the fecal samples. **c.** The metabolic output of the characteristic strain for resistant dextrin (RD), *Acutalibacer* sp. only partially matched that of the average fecal community. The lower panel shows the pairwise Spearman correlations of the measured metabolites across the RD enrichments with a strong anticorrelation between SU and PR. Because SU is typically converted only to PR they are combined for a better match to the profile of the characteristic taxon. **d.** Enrichments in basal medium supplemented with SU reveal a binary pattern of SU consumption by the fecal microbiota. Positive values show the concentration of produced metabolites and negative values consumed metabolites. **e.** Three genera were identified as characteristic for SU consumption, with a strong binary pattern across the five microbiota for which SU consumption was observed. **f.** The heatmap shows the relative abundances of each of the characteristic genera in the eight fecal samples. The presence or absence of the three genera combined predicts in which microbiota SU accumulates in enrichments on primary substrates. **g.** Metabolic output of enrichments in basal medium supplemented with lactate (LT). **h.** Four genera were identified as characteristic for LT consumption, with only *Anaerobutyricum* and *Coprococcus_A* widely present across donor microbiota. **i.** *Coprococcus_A* and *Frisingicoccus* show signs of a putative guild for lactate-to-propionate conversion.

The metabolites produced by the characteristic taxa matched those produced by the whole fecal microbiota. In PE, *L. eligens* produced acetate (26.6 mM carbon) and formate (9.0 mM carbon) at a molar carbon ratio of 3:1. This was consistent with what was observed across all the fecal microbiota (Figure 3b; mean acetate = 30.2 mM carbon, formate = 8.0 mM carbon, approx. molar carbon ratio 3.77:1). In RD, *Acutalibacter* sp. produced acetate (9.2 mM carbon) and succinate (13.9 mM carbon) at a molar carbon ratio of 2:3 (Figure 3c). This did not match the metabolite profiles across donors, where propionate and butyrate was also produced (Figure 3c). Propionate production was bimodal across donors, with 10-14 mM carbon in some (M1, M3, M7, M8) and 0-4 mM carbon in the others (M2, M4, M5, M6). Succinate concentrations were strongly anticorrelated with propionate along same bimodal partitioning of donors (Figure 3c). Because propionate is the typical fermentation product of the succinate pathway^22^, we hypothesized that this dichotomy was the result of incomplete succinate utilization in those four microbiota for which succinate was measured. To test this, we performed additional enrichments using succinate as the supplemented carbon source.

Enrichments of the fecal microbiota in succinatesupplemented medium (SU) also had a binary pattern of succinate consumption and propionate production. Succinate was consumed by five of the eight donor microbiota (M1, M3, M6, M7, M8) and was converted to propionate at a molar ratio of 1:1, but was not consumed at all by the remaining three microbiota (Figure 3d). This binary pattern of succinate consumption/propionate production almost perfectly matched the binary pattern we observed on RD, confirming that the accumulation of succinate was due to the inability of those donor microbiota to consume the succinate produced from RD.

### Strong guild structures arise at the secondary consumer level

To further understand the dichotomy of succinate consumption on a taxonomic level, we determined the characteristic taxa for succinate using an equivalent procedure as for the primary substrates. After filtering for unspecific growth, we identified only three characteristic genera for SU consumption (Figure 3e, Supplementary Figure S9), all of which belong to the *Negativicutes* class that is known for the ability to consume succinate^22^. Within donor microbiota, only a single characteristic genus was respectively enriched—or none at all: *Phascolarctobacterium* in M1 and M3, *Phascolarctobacterium_A* in M7 and M8, and UBA1822 (*Dialisteraceae* family) in M6. Such strong mutual exclusivity is a hallmark of functional guilds, and we thus postulated that these three genera form a functional guild for SU consumption. In this case, the abundance of the functional guild as a whole in the diluted fecal sample should determine whether SU accumulates or is consumed.

The mutual exclusivity of the three genera was mirrored in the composition of the fecal microbiota themselves (Figure 3f). Three of the microbiota did not have detectable levels of any of the SU guild members in the fecal samples (M2, M4, M5) and these three microbiota generally accumulated SU across all supplemented substrates (Figure 3f). In contrast, the other five fecal microbiota had substantial abundances of the SU guild and did not accumulate SU in the enrichments. Thus, the abundance of the SU guild in the fecal samples can explain why succinate accumulates in certain fecal microbiota while in others it does not.

Having observed the prominent guild structure for SU consumption we asked whether this was also true for the other intermediate metabolites, and performed equivalent enrichments for formate (FO) and lactate (LT). FO was not consumed during the 48 h cultures in these specific culture conditions. In contrast, LT was fully consumed by all eight microbiota and was primarily converted to butyrate (Figure 3g). We did, however, observe differential propionate production from LT across donor microbiota, with 3-6 mM produced in five microbiota (M1, M2, M4, M7, M8) and none produced in the remaining three (M3, M5, M6). We identified four characteristic genera for LT consumption, of which *Anaerobutyricum* and *Coprococcus_A* were selected for across almost all microbiota (8/8 and 7/8, respectively). These two genera are known LT consumers but use different pathways, where *Anaerobutyricum* (e.g. *A. hallii*) produces butyrate and *Coprococcus_A* (e.g. *C. catus*) produces propionate^23, 24^. The more strongly that *Coprococcus_A* was selected for, the more propionate was produced (Spearman’s *r* = 0.786, *p* = 0.048), suggesting that *Coprococcus_A* is a key driver of propionate production from LT. Inspired by the SU guild, we hypothesized that *Anaerobutyricum* and *Coprococcus_A* would directly compete for LT. However, the *A* were not anti-correlated (Spearman’s *r* = 0.551, *p* = 0.157) suggesting some mechanism for coexistence of these two characteristic genera. In M4, *Coprococcus_A* was absent and instead *Frisingicoccus* was selected for (Figure 3i). *Frisingicoccus* is a direct phylogenetic neighbor of *Coprococcus_A*^25^, such that it likely uses the same metabolic pathways for LT which could explain the observed propionate production in M4 without *Coprococcus_A*. Thus, while guilds are not as evident at first glance for LT as for SU, a guild-like structure does emerge among those LT consumers that employ similar metabolic pathways.

### A nutritional intervention study confirms the characteristic taxa for resistant dextrin

The characteristic taxa provide predictions of which bacteria are stimulated by different components of human diet. To test these predictions, we made use of microbiome data from a placebocontrolled randomized nutritional intervention study using RD. Briefly, individuals were given either RD (NUTRIOSE®) or maltodextrin—a placebo that is completely metabolized in the small intestine (Figure 4a). Fecal samples were collected prior to the start of the study and after four weeks of dietary supplementation, and their composition was analyzed using 16S amplicon sequencing (see Methods).

**Figure 4:**
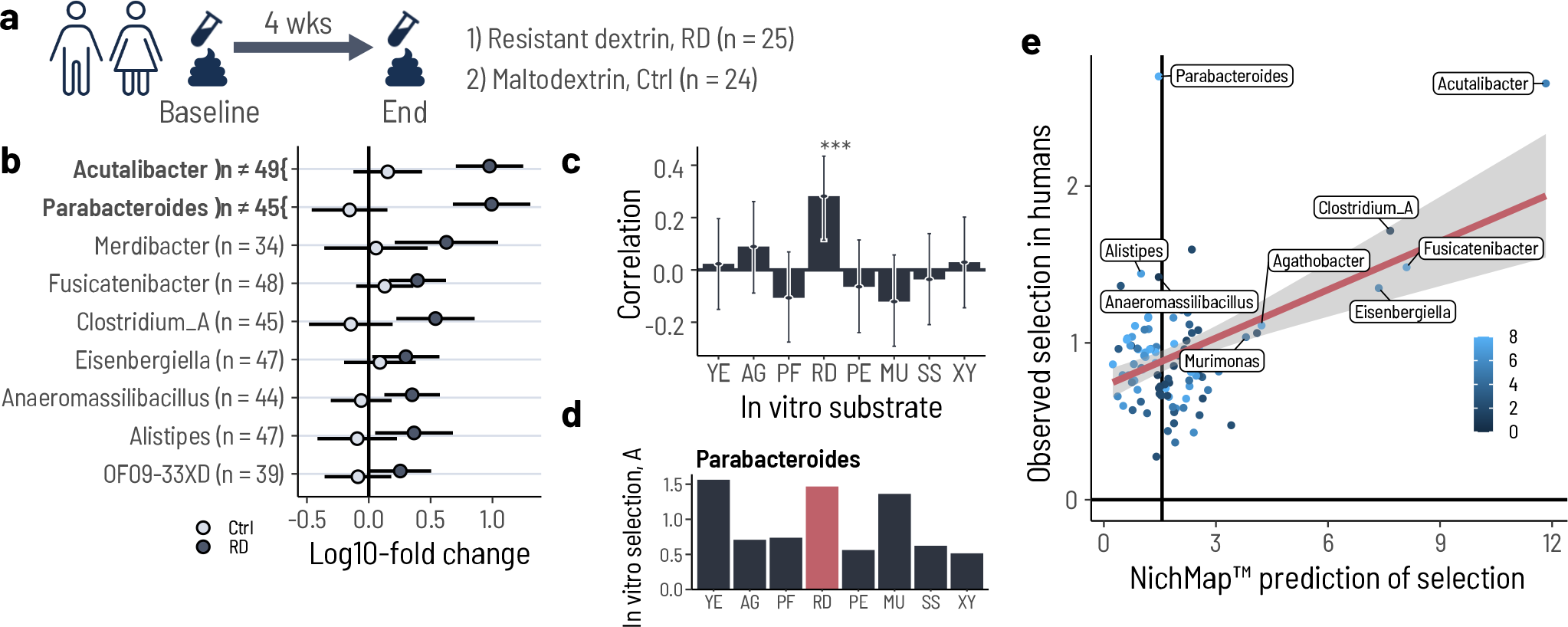
Microbiota changes in a nutritional intervention study with resistant dextrin recapitulate the predictions from the NicheMap™. **a.** Healthy participants in the study supplemented their otherwise normal diet daily for 4 weeks with either NUTRIOSE®(resistant dextrin; RD) or maltodextrin as a control (Ctrl). Fecal samples were collected at the start and end of the study and analyzed with 16S amplicon sequencing. **b.** Only few genera were differentially changed during the study in the RD group compared to the Ctrl group. Genera that increased significantly in relative abundance in the RD group from start to end of the study (dark circles). The corresponding change in the Ctrl group is shown with light circles. **c.** Of all enrichment conditions, only RD correlated significantly with the changes observed in the nutritional intervention study. The bars show the Pearson correlation between in vitro enrichment and in vivo change in relative abundance. The error bars for the estimated confidence interval for the correlation coefficient. **d.** The overall magnitude across all substrates of Parabacteroides enrichment *in vitro* is low, but is most strong on RD compared to the other dietary fibers and sugars. **e.** The magnitude of selection on the NicheMap™ of individual genera predicts the change in relative abundance in the study.

Four weeks of supplementation with RD resulted in a larger change in the microbiota of study participants compared to supplementation with the placebo (Supplementary Figure S10). However, the overall microbiota diversity did not change significantly in either group. Both these results are consistent with those from a similar nutritional intervention study with NUTRIOSE®^26^. Only two genera, *Acutalibacter* and *Parabacteroides*, significantly increased over the four weeks in the RD group compared to the placebo group (Figure 4b), with *Parabacteroides* and *Acutalibacter* increasing from a mean relative abundance of 0.5% and 0.2% at baseline to 3.4% and 3.0% in the RD group at the end of the study, respectively (Supplementary Figure S11). Additional genera increased by a degree that was significantly higher than zero, but not significantly different from the placebo group in the statistical sense: *Merdibacter*, *Fusicatenibacter*, *Clostridium_A*, and *Eisenbergiella* (Figure 4c).

The genera that increased in the microbiota of study participants after four weeks of RD supplementation was in good agreement with the characteristic taxa of RD from the *in vitro* enrichments. We compared the degree of selection for genera measured *in vitro* to the change in relative abundance in the study participants over the four weeks of RD supplementation. To avoid comparing taxa that were not present in the donor microbiota, we computed the expected number of donor inocula out of the eight that contained a genus of interest, *p*_2_, and only included those with *p*_2_ *>* 1. RD was the only *in vitro* substrate for which the estimated degree of selection correlated significantly with the changes in the patients (Figure 4c; Pearson’s *r* = 0.45, *p* = 9.92 *×* 10*^−^*^7^). Almost all of the characteristic taxa identified *in vitro* were also selected for in the human participants during the course of the study (Figure 4e). The exception was *Parabacteroides*, which was strongly selected for in the study participants, but was not among the characteristic genera for RD. This was because the overall selection for *Parabactoides in vitro* was low in magnitude but was nevertheless strongest for RD compared to all the other tested dietary fibers (Figure 4d). This suggests that the predictions of selection from the NicheMap™ are specific for those taxa that supplementation with RD (i.e. NUTRIOSE®) selects for in humans.

## Discussion

A major impediment to manipulating and engineering gut microbiomes is the lack of a good catalog that maps the different microbial constituents of intestinal bacteria to their respective functional roles. Inspired by approaches from chemical engineering, we and others have put forward blueprints of how microbiomes might be organized and reconstructed based on a desired metabolic output^13, 27, 28^. Yet, without a systematic mapping of bacteria onto the different roles that make up these blueprints, such approaches retain an *ad hoc* nature. Most current efforts to generate such a map take a genomics-based approach, fueled by the commoditization of highthroughput sequencing and better computational and statistical tools. Extrapolating from metagenomes, metatranscriptomes, or other meta-omes to actual phenotypes is challenging^29^, and further projecting this into a community context is typically unfeasible^30^.

The approach we present here circumvents many of these obstacles by directly measuring bacterial metabolic phenotypes in their relevant complex community context. We demonstrated the power of our approach by identifying the characteristic bacteria that are associated with specific parts of intestinal carbohydrate fermentation. The characteristic taxa we identified comprised both expected taxa as well as novel taxa. For example, the *Lachnospira* we identified as characteristic for pectin are known to be “pectinophilic”^31^, but we were also able to map the more elusive genera UMGS1441 and CAG-41 to pectin degradation. CAG-41 is an unclassified *Firmicutes* bacterium that was first identified in MetaHIT^32^, and since then has been repeatedly reported as differentially abundant, e.g. between health and disease^33–35^. Similary, UMGS1441 has most recently been assembled from metagenomic data from chickens (*Candidatus* Gallispira)^36^. For AG, the top characteristic genera included an unclassified genus of *Lachnospiraceae* (here Lachnospiraceae_Genus) and UMGS1375. Lachnospiraceae_Genus grouped together ASVs whose closest BLAST hit was *L. eligens* at < 95 % identity. The ASVs that grouped into UMGS1375 were a close match (97 %-99 % identity) to *Hominisplanchenecus faecis*, a genus that was only very recently isolated using an automated high-throughput approach^37^ but not yet characterized. The characteristic genera for RD, *Acutalibacter* and *Eisenbergiella*, have been observed in gut microbiomes^38, 39^ though their role has remained unclear so far, and *Clostridium_A* (e.g. *C. leptum*/Cluster IV) has been a group of interest for over a decade^32, 40^. The NicheMap™ thus offers an approach to phenotypically attribute function to these ubiquitous yet elusive gut bacteria.

We were thus able to map an important part of key intestinal bacteria across human fecal microbiomes onto the biochemical scaffold for carbohydrate fermentation—the assumption being that this is a good representation of the key metabolic role of the microbiome. Doing so allowed us to attribute the concepts of context-dependence and functional guilds from ecological theory to the observed structure and further refine the functional blueprint for gut microbiomes. The premise is that intestinal microbiomes are organized in a trophic hierarchy, with different organizational structures at each trophic level (Figure 5). ‘Complex primary degraders’ break down chemically complex energy sources, such as dietary fibers, often doing so extracellularly and thus releasing the simple building blocks to the local environment. On the next level, the resulting simple sugars are further metabolized by ‘simple primary degraders’, into either fermentation intermediates or end products. Because the final steps of the fermentation cascade typically provide less energy to the microbe, intermediate products—in particular, formate, lactate, and succinate—are often released back into the environment. At the bottom level, secondary consumers are specialized in the conversion of these fermentation intermediates into fermentation end products, mostly short-chain fatty acids. While this concept of trophic units is not new *per se*^16, 18, 41^, we here develop an understanding of how these trophic units are organized. Each of the trophic units have a unique combination of taxonomic diversity, guild structure, and niche diversity. The taxonomic diversity within a niche follows a ‘bell shape’, with few taxa associated with complex primary substrates and secondary consumers, and higher diversity in the simple substrate consumers (Figure 5). At the top trophic level, complex carbohydrates with high heterogeneity in chemical structure require a repertoire of various enzymes to break down the different kinds of glycosidic bonds. At the bottom trophic level, the metabolic reactions are often rather specific in terms of the physiochemical environment (pH, redox, etc.) and the energetic yields are low. Both extrinsic constraints impose a high degree of evolutionary selection that limits diversification. At the middle level, the main metabolic processes revolve around the common fermentative pathways that are broadly distributed.

**Figure 5:**
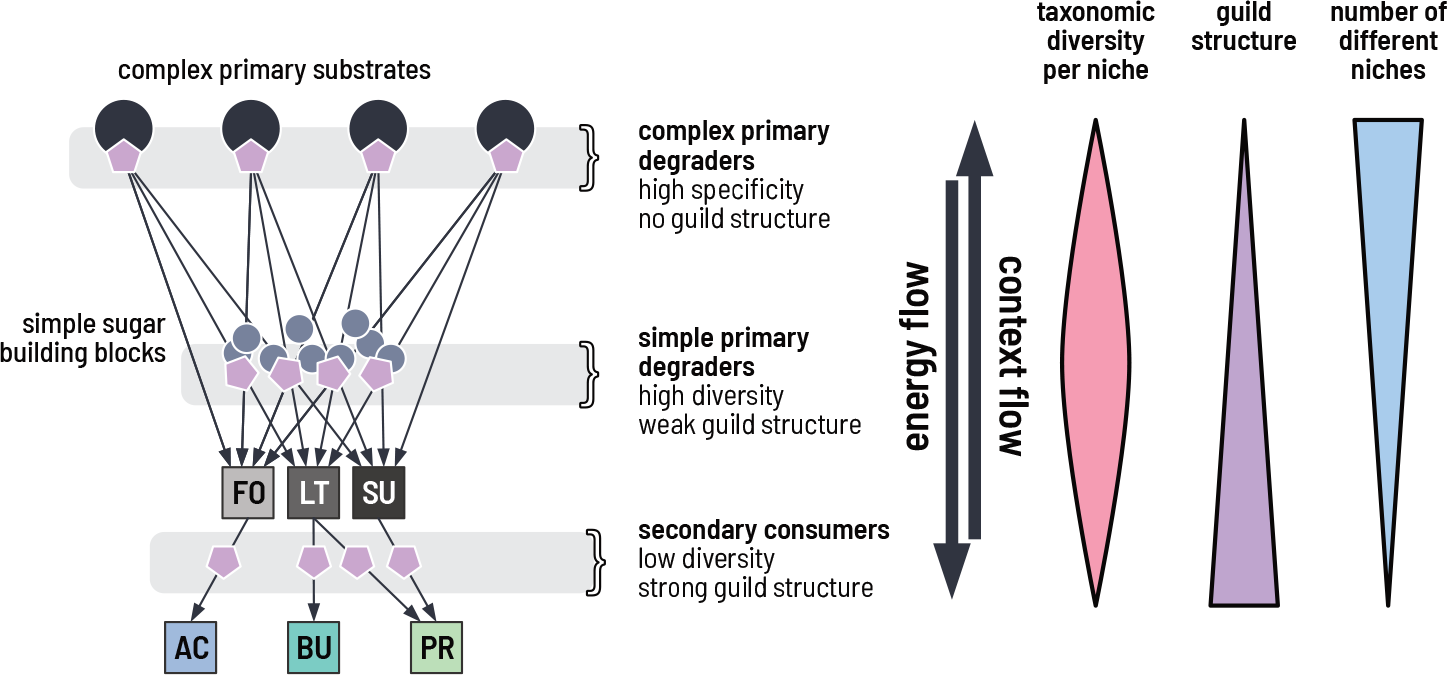
Microbiomes are organized into niches in a hierarchical manner that follow the scaffold of thermodynamic energy flow. Microbiome taxa can be mapped onto a metabolic reaction scaffold with three trophic levels. Complex primary degraders are at the top level and break down complex energy sources like dietary fibers. The mapping of taxon-to-fiber is highly specific. At the middle level are simple sugar/substrate utilizers. The phylogenetic diversity at this level is high with some guild-like properties. Finally, at the bottom level are secondary consumers. The mapping of taxon-to-niche is also highly specific but with some redundancy that manifests is a strong guild structure. Energy flow and accessibility is determined top-down, while the context that directs the metabolic output is determined bottom-up.

Guild structure is weakest at the complex primary degrader level. The breadth of substrate types combined with the specialization required to degrade a particular substrate result in few taxa that sufficiently overlap in terms of their substrate repertoire to lead to competitive exclusion. Examples for this in our data is the conservation of the CAG-41 and *Lachnospira* for PE, and *Acutalibacter* for RD. At the simple sugar level, diversity is too high to allow for a clean guild structure to emerge— a multitude of gut bacteria can utilize sugar monomers (glucose, galactose, xylose, etc.) as an energy source. Finally, at the secondary consumer level, guilds are very pronounced, possibly also because the low energetic yield increases the perceived competition for energy. We observed that typically only a single succinate consumer was dominant per fecal microbiome, and that the function ‘broke down’ if the whole guild was absent. Surprisingly, for lactate we observed coexistence of at least two types of lactate consumers, *Anaerobutyricum* and *Coprococcus_A*/*Frisingicoccus*, despite supposedly strong competition for lactate. These lactate consumers differ in the metabolic pathway they encode, with the former producing butyrate and the latter propionate^24^. A strong guild structure possibly applies within the lactate-to-propionate consumers (*Coprococcus_A* and *Frisingicoccus*). We thus propose that niches—and thus the functional guilds that associate with them—should be defined taking into account both inputs and outputs, and the physiochemical parameters that modulate these inputs and outputs.

Finally, the variety of different niches at each trophic level decreases from top to bottom. The combination of all three aspects leads to the predictions of different amounts of taxonomic diversity at each trophic level. First, at the complex degrader level, overall taxonomic diversity is high, but this is driven by the diversity in substrates. Second, at the simple degrader level, taxonomic diversity is also high, but here this is driven by the competition for a small variety of simple sugars. Third, at the secondary consumer level, taxonomic diversity is low, driven by the combination of very low niche diversity and low energetic yields. Taken together, this implies opposing forces that drive gut microbiome structure and function. Energy flows ‘top-down’, determined first by the available substrates and their corresponding degraders. Conversely, context flows ‘bottom-up’, with the guild composition at the secondary degrader level ultimately determining the metabolic fate of the degraded carbon sources and thus the overall community output.

Our approach complements other methods to describe how gut microbiome function is structured^42^. Importantly, it provides much needed quantitative biological data to inform computational models^43^. As a consequence of measuring the biological phenotype directly using low-throughput microbiological experiments, instead of high-throughput genomic approaches, our study is limited in terms of the number of different microbiomes we screened. Nevertheless, by choosing these along the enterotype gradient^44^, we managed to obtain a representative sample of the described microbiome diversity. Our competitive growth assay also imposes a strong bottleneck in terms of initial dilution of the fecal sample which we account for using a careful probabilistic model. By design, this puts a focus on those taxa that have sufficient abundance to drive the metabolic conversions in the gut. Despite the small number of fecal microbiota and initial bottleneck, we were able to make predictions that were validated in a nutritional intervention study. Such studies typically pose a challenge because lifestyle factors including the remainder of the individual’s diet is not controlled. Nevertheless, we managed to capture most of the *in vivo* microbiome response. This is possible because of the high degree of conservation at the complex degrader level. We therefore expect our approach to also be accurate in the prediction of the microbiome response to other dietary fibers or nutritional supplements, allowing for fiber design and pre-screening prior to costly intervention studies and coming a step closer towards personalized nutrition^45^.

Overall, we have put forward an approach to generate a structure-function map for gut microbiomes that is based on context-aware phenotypic data. This has the important advantage over genomic or statistical methods by directly measuring what functions are performed by a complex microbiome and identifying putative bacteria that are phenotypically associated with this function. Having such a map on hand will contribute to a better understanding of microbiome function and dysbiosis, thus enabling better targeted functional interventions.

## Acknowledgements

We thank Shaul Pollak for feedback on the manuscript.

## Data availability

The data that support the findings of this study are provided as supplementary files for peer review and will be available from the corresponding author upon reasonable request.

## Author Contributions

L.A., C.M., and M.R. performed experiments. T.d.W., L.A., C.T., C.L., O.X.C., and G.E.L. planned the execution of the project. L.A., P.R.v.B., and G.E.L. analyzed the data. T.d.W., B.R., and G.E.L. conceived the project. C.T. conceived and oversaw the nutritional intervention study. M.M. identified the donors based on enterotype. L.A., P.R.v.B., C.L., C.T., T.d.W., and G.E.L. interpreted the results. G.E.L. and L.A. wrote the first draft of the manuscript. All authors reviewed the final manuscript and provided feedback.

## Methods

### Collection of feces

The research project was approved by the Ethic Committee of the Canton of Zurich (2017-01290). Fresh fecal samples were donated from eight healthy individuals with no history of antibiotic use, intestinal infections, or severe diarrhea during the three months prior to making the donation. The donors did not take immunosuppressive drugs, blood thinners, or medication affecting the bowel passage or digestion. Fecal samples were anaerobically transported in an airtight container together with an Oxoid™ AnaeroGen™ 2.5 L sachet (Thermo Fisher Diagnostics AG, Pratteln, Switzerland) and processed within three hours after defecation. Stool consistency was evaluated optically according to the Bristol Stool Scale^46^ and samples within the defined range of a healthy stool, notably with a score between 3-5, were accepted.

### Culture media and anaerobic dilution solution

Culture media were based on a common basal medium (Supplementary Table S1) and supplemented with nine distinct growth substrates (Figure 1a). All medium ingredients except sodium bicarbonate and L-cysteine HCl were dissolved in an Erlenmeyer flask and the pH was adjusted to 7 by titrating 5 mM sodium hydroxide. The media were boiled for 15 min for major removal of oxygen, under constant moderate stirring, and using a Liebig condenser to prevent vaporization of ingredients. After boiling, the media were constantly flushed with CO_2_. Sodium bicarbonate and L-cysteine hydrochloride monohydrate were added when the media cooled down to 55 *^◦^*C for further reduction of residual oxygen for 10 min. Aliquots of 8 mL of medium were filled into Hungate tubes under constant flushing with CO_2_, and Hungate tubes were sealed with butyl rubber stoppers and screw caps (Millan SA, Geneva, Switzerland). The media were sterilized by autoclaving and subsequently stored at room temperature. Anaerobic dilution solution for fecal samples (Supplementary Table S2) was prepared following the same procedure as for the culture media, except that aliquots of 9 mL were filled into Hungate tubes to facilitate serial dilutions.

### Feces processing and dilutions

For processing, the fecal samples were transferred into a Coy anaerobic chamber (Coy Laboratories, Ann Arbor, MI, USA) with an atmosphere of 10 % CO_2_, 5 % H_2_, and 85 % N_2_. We prepared a 1:10 dilution with approximately 1 g of fecal sample that was measured with a sterile plastic spoon (VWR International, Dietikon, Switzerland) and subsequently suspended in 9 mL of anaerobic dilution solution. We transferred 1 mL of the 1:10 dilution into 9 ml of anaerobic dilution solution to obtain a 1:100 dilution. We then transferred 1 mL of the 1:100 dilution into a sterile Hungate tube containing 9 mL of anaerobic dilution solution. We performed subsequent serial dilutions in steps of 10 down to 10*^−^*^1^^1^ outside of the anaerobic chamber under sterile, anaerobic conditions using the Hungate technique^47^.

### *In vitro* enrichments

Anaerobic *in vitro* enrichments were performed in Hungate tubes sealed with butyl rubber stoppers and screw caps (Millan SA, Geneva, Switzerland). For each fermentation, 0.3 mL of the 10*^−^*^8^ fecal sample dilution was inoculated into 8 mL (ca. 10*^−^*^9^ dilution) of cultivation medium under sterile and anaerobic conditions using the Hungate technique. All cultures were incubated at 37 *^◦^*C. After 48 h of incubation, we measured optical density at a wavelength of 600 nm (OD600) directly in the Hungate tubes with a WPA CO 8000 Cell Density Meter (Biochrom Ltd, Cambridge, England).

### Microbial metabolite analysis

Metabolite concentrations of acetate, propionate, butyrate, lactate, succinate, formate, and ethanol were measured by HPLC analysis. Samples were prepared from 1 mL of bacterial culture centrifuged at 14 000 g for 10 min at 4 *^◦^*C. The supernatant was filtered into 2 mL short thread vials with crimp caps (VWR International GmbH, Schlieren, Switzerland) using non-sterile 0.2 µm regenerated cellulose membrane filters (Phenomenex Inc., Aschaffenburg, Germany). A volume of 40 µL of sample was injected into the HPLC with a flow rate of 0.6 mL*/*min at a constant column temperature of 80 *^◦^*C and using a mixture of H_2_SO_4_ (10 mM) and Na-azide (0.05 g*/*L) as eluent. Analyses were performed with a Hitachi Chromaster 5450 RI-Detector (VWR International GmbH, Schlieren, Switzerland) using a Rezex ROA-Organic Acid (4 %) precolumn connected to a Rezex ROA-Organic Acid (8 %) column, equipped with a Security Guard Carbo-H cartridge (4 *×* 3 mm). Metabolite concentrations were determined using external standards (all purchased from Sigma-Aldrich, Buchs, Switzerland) via comparison of the retention times. Peaks were integrated using the EZChromElite software (Version V3.3.2.SP2, Hitachi High Tech Science Corporation).

### DNA extraction

For fecal samples, we extracted total genomic DNA from 200 mg of each sample. For cultures, we centrifuged 1 mL of bacterial cultures at 14 000 g and 4 *^◦^*C for 10 min. For both sample types, we used the FastDNA®SPIN Kit for Soil (MP Biomedicals, Illkirch Cedex, France) according to the manufacturer’s instructions. We quantified the total DNA concentration using the Qubit®dsDNA HS Assay kit (Thermo Fisher Scientific, Pratteln, Switzerland).

### Amplicon sequence variants and taxonomic assignment

We performed amplicon sequencing of the 16S rRNA V3-V4 region on the MiSeq platform (Illumina, CA, USA) using the primer combination 341F (5’-CCTACGGGNBGCASCAG-3’) and 806bR (5’-GGACTACNVGGGTWTCTAAT-3’). Library preparation and sequencing was performed by StarSEQ GmbH (Mainz, Germany) with 25 % PhiX to balance the composition of bases. Amplicon Sequence Variants (ASVs) were inferred using Dada2 v1.18.0^48^ with read length filtering set to (250, 210), maxEE set to (4,5), inference done in ‘pseudo pool’ mode. Read pairs were merged with minimum overlap of 20, and bimeras were removed using the ‘consensus’ method. Taxonomic assignment was performed with the assignTaxonomy function from Dada2 using GTDB r95 prepared for Dada2^49^. In order to retain all ASVs but nevertheless be able to interrogate composition at different taxonomic levels, we used a bootstrap cutoff of 0.2 for each taxonomic level and propagated the assignment using the lowest well-assigned taxonomic level: for example, Lachnospiraceae_Genus groups together all ASVs that have a clear family-level assignment, *Lachnospiraceae*, but no genus-level assignment. For some of these ASVs bootstrap values were low because of the high degree of degeneracy of reference sequences (e.g. *Bifidobacterium* sp. and *Escherichia* sp.). For these ASVs, we further refined the assignment by aligning to the reference GTDB database manually.

### Quantification of total viable cells in feces

We estimated the total number of viable cells in feces by most probable number (MPN) enumeration in liquid culture and using strict anaerobic Hungate techniques. To this end, for each fecal sample we inoculated 0.3 mL of the 10*^−^*^9^, 10*^−^*^10^, and 10*^−^*^11^ dilutions into 8 mL of M2GSC growth medium in triplicates. We categorized positive growth in a tube as an OD600 above 0.5. We performed a Bayesian estimation of the concentration of viable cells, *λ*, in the fecal samples by fitting a binomial model to the number of tubes for which we observed growth. We used a Gamma prior on *λ* with *α* = 1 and *β* = 0.01 and sampled from the posterior with RStan^50^. All samples had viable cell numbers within a range of 10^10^ *−*10^12^ cells per gram of feces (Supplementary Figure S11), as expected for healthy stool^5^^1^.

### Estimation of cell concentrations from total DNA

We fit a statistical model to predict cell densities [cells/ml] from the concentration of total extracted DNA [ng/ml]. To this end we used data from pilot enrichments that were performed on the same panel of carbon sources apart from AG and YE, and using the same methodology with substrates added at a concentration of 6 g*/*L. The remainder of the protocol was identical to what is described. For each of the enrichment cultures, we measured absolute abundances using two methods: (i) the concentration of total extracted DNA using the Qubit®dsDNA HS Assay kit; (ii) total bacterial cell concentrations with qPCR using universal 16S primers. We then fit a linear model relating the measured log DNA concentration, *x*, to the measured log cell concentration, *y*. Because both *x* and *y* are measured quantities of a true value *x*_0_ and *y*_0_ for *x* and *y*, respectively, we used a Bayesian approach where *x ∼ N* (*x*_0_*, σ_x_*), *y ∼ N* (*y*_0_*, σ_y_*), and *y*_0_ = *a* + *bx*_0_. We coded this model in Stan with priors *σ_x_*, *σ_y_ ∼* Γ(0.1, 0.1), *a ∼ N* (0, 5), *b ∼ N* (0, 1), and *x*_0_ *∼ N* (0, 10). Posterior samples (*n* = 4000) were generated with RStan. Sampling from this full model resulted in a strong anti-correlation between *a* and *b*, and a mean posterior estimate of *b* = 0.98 very close to 1. Because we expect the DNA concentration and cell concentration to scale proportionally on a linear scale (*b* = 1), we repeated the fit with *b* fixed to 1. This resolved the posterior parameter correlations and hence we used this simplified method to convert DNA concentration to cell concentrations in this setup (Supplementary Figure S12).

### Estimation of total carbon extraction and total bacterial growth

We performed Bayesian estimation of the mean total carbon extracted from the primary substrates. We first estimated the total concentration of carbon in each sample by multiplying the individual measured metabolite concentrations by the respective carbon count of the molecule and summing over all measured metabolites. Note, that this estimation is a lower bound on the actual total carbon, as it only uses the measured metabolites and does not account for e.g. CO_2_. We then fit a model with a joint baseline, *ξ_d_*, across all growth conditions including the basal medium (mM2) without added substrate, a mean effect, *ϕ_s_*, for each substrate, and a donor-substrate specific deviation, *ψ_d,s_*. This has the advantage of using all of the data to estimate *ξ_d_* and is more conservative in terms of uncertainty than using growth on mM2 alone. We used very weakly normally distributed priors for *ξ ∼ N* (30, 50) and *ϕ ∼ Normal*(50, 30) that were roughly centered around the observed concentrations from the pilot experiment to improve seeding of the Markov chain Monte-Carlo (MCMC) chains. We performed the Bayesian equivalent of LASSO regularization using Laplace priors for the *ξ_d,s_* with a hyperparameter *σ_ψ_* for the variance, *ψ_d,s_ ∼* Laplace(0*, σ_ψ_*) and *σ_ψ_ ∼* InvGamma(0.1, 0.1). We modelled the residual error across replicates as *ε ∼ N* (0*, σ_ε_*), where *σ_ε_ ∼* InvGamma(0.1, 0.1). We coded the model in the Stan language and performed posterior sampling using RStan. We used the same model to estimate the total amount of bacterial growth, just using the estimated log_10_ cell concentrations instead of the total carbon concentration.

### Computation of relative enrichment

We first computed the expected relative abundance in a sample, *x_i_* = (*n_i_* + *j_i_*)*/N*, where *n_i_* is the number of sequencing reads, *j_i_* = 1 if the taxon was observed in any of the enrichments inoculated with that specific fecal sample and *j_i_* = 0 otherwise, and *N* is the total number of sequencing reads in the sample. The Bayesian interpretation of the pseudo count *j_i_* = 1 corresponds to computing the posterior mean under the assumption of a multinomial sampling model with a flat Dirichlet prior for those genera that were observed at least once in at least one sample. We computed the absolute abundances, *X_i_*, by multiplying *x_i_* with the estimated cell concentrations for each sample.

To account for the potential loss of a taxon *i* during the dilution at the inoculation step, we computed the probability that a taxon is inoculated with at least one cell in a culture. We used the posterior samples of the fecal cell densities, *λ*, using the MPN method, and assumed that the total number of cells, *M*, that are retained in the inoculum that is diluted *ϑ*-fold is sampled from a Poisson distribution with rate *λϑ*. The number of cells of each taxon, *m_i_*, that make it into the inoculum is then sampled with *M* draws from a multinomial distribution with probabilities equal to the relative abundance in the fecal sample, *z_i_*, where *z_i_ ∼* Dirichlet(*n*+*j*), equivalent to the computation of *x_i_* above. Using the posterior samples of *m_i_*, we compute the probability of loss to dilution, *p*_0_*_,i_*, as the fraction of posterior draws that give *m_i_* = 0. Furthermore, we define *p*_1_*_,i_* as the relative abundance of taxon *i* in the inoculum sample conditioned on not having been lost.

We then computed a weighted average for the estimated relative and absolute abundances over the triplicates that accounts for potential loss to dilution in the inoculum, *y_i_* = (*x_i_*_1_*w_i_*_1_ + *x_i_*_2_*w_i_*_2_ + *x_i_*_3_*w_i_*_3_)*/*3, where *w_ik_* = 1 if the taxon was observed in the *k*-th replicate of the enrichment culture, and *w_ik_* = 1 *− p*_0_*_,i_* if it was not observed. We then computed the relative enrichment of a taxon in a specific condition as the change in log_10_ relative abundance from a reference composition, *E_i_* = log_10_ *x_i_ −* log_10_ *x*^(ref)^, where the reference was typically the diluted fecal composition, *p*_1_, or the enriched composition, *y*, in mM2. We also computed the absolute enrichment, *A_i_* = log_2_ *X_i_ −* log_2_ *X*^(ref)^, and the biomass increase *B_i_* = *X_i_* − *X_i_*^(ref)^.

### Nutritional supplementation study with NUTRIOSE®

The study protocol and annexes were submitted to the CPP II Île-de-France Ethics Committee (Saints Péres) on April 18, 2012. The project was analyzed on May 3, 2012 and additional information was requested. The complementary file was analyzed by the ethics committee on May 22, 2012 and final positive opinion was given on May 24, 2012. The AFSSAPS identifier (ID RCB) is 2012-A00061-40. The protocol was registered on ClinicalTrials.gov (NCT01897649). The study was a monocentric, randomized, controlled, and double-blind study, conducted with two parallel groups: Control group (or comparator) and Test group.

The intervention phase of the study lasted 35 days from the date of inclusion in the study. Sixty subjects were recruited during the pre-selection visit and included in the protocol during the inclusion visit. Subjects were randomized in two groups (thirty subjects per group). Each subject consumed the tested product (15 g NUTRIOSE®FB06) or the comparator (GLUCIDEX®21) during 28 days from day 1 (D1) to day 28 (D28). Subjects were asked to give a stool sample at the inclusion visit one week before D1 (D-7), 14 days after the beginning of product consumption (D14), and at the end of the study (D29). Subjects were asked to record food intake for three non-consecutive days including two weekdays and one weekend day the week before D1, D15, and D28.

NUTRIOSE®FB06 is a soluble dietary fiber produced from wheat starch, manufactured, and marketed by Roquette Fréres Company. NUTRIOSE®was provided in the form of a white powder, odorless, tasteless. The product was packaged in aluminum sealed bags. Each bag contained 15 g NUTRIOSE®FB06. GLUCIDEX®21 is dried glucose syrup derived from corn starch. It is composed of 3 % glucose, 7 % maltose and 90 % of glucose polymers with polymerization degree larger than 3. GLUCIDEX®21 is in the form of a white powder with a slightly sweet flavor. Each subject consumed the content of one bag (NUTRIOSE®or comparator product) per day during 28 days.

Stool samples were collected at D-7, D14, and D29. Defecation took place at home on the morning of the medical visit or at the research center. A thermal protector device allowing temperature maintenance at 4 *^◦^*C for 12 h was provided to volunteers for stool transportation from home to hospital. Stools were kept at 4 *^◦^*C at the research center until sampling and being frozen at *−*80 *^◦^*C. The transport of frozen stool samples at *−*80 *^◦^*C to the laboratory in charge of microbiological analysis was performed using an adequate volume of dry ice. A 25 mg lyophilised sample of feces was used for extraction of total bacterial DNA using a commercial kit (QIAamp DNA stool kit; Qiagen). 16S amplicon sequencing of the V3-V4 region was performed on Illumina MiSeq (2 *×* 300 bp) at DNAVision (Gosselies, Belgium). ASVs were inferred using the same pipeline as described for the enrichments.

## Supplementary Tables

**Table S1:** Base medium (mM2) and anaerobic dilution solution (ADS) composition

| Product | mM2 | ADS | Supplier | Article No. |
| --- | --- | --- | --- | --- |
| Amicase | 1 g/L |  | Sigma-Aldrich Chemie GmbH, Buchs, Switzerland | A2427 |
| Yeast extract | 1.25 g/L |  | Sigma-Aldrich Chemie GmbH, Buchs, Switzerland | 1.11926 |
| Meat extract | 0.5 g/L |  | Sigma-Aldrich Chemie GmbH, Buchs, Switzerland | 70164 |
| Rumen fluid | 300 mL/L |  | Schlachthof Zurich |  |
| Sodium bicarbonate | 4 g/L | 4 g/L | Sigma-Aldrich Chemie GmbH, Buchs, Switzerland | 13433 |
| Potassium phosphate dibasic trihydrate | 0.45 g/L | 0.59 g/L | Sigma-Aldrich Chemie GmbH, Buchs, Switzerland | P5504 |
| Potassium dihydrogen phosphate | 0.45 g/L | 0.87 g/L | VWR International AG, Dietikon, Switzerland | 26923.298 |
| Sodium chloride | 0.9 g/L | 1.74 g/L | Thermo Fischer Scientific, Pratteln, Switzerland | AC44730 |
| Ammonium sulfate | 0.9 g/L | 1.74 g/L | Sigma-Aldrich Chemie GmbH, Buchs, Switzerland | A5132 |
| Magnesium sulfate water free | 0.09 g/L | 0.17 g/L | Thermo Fischer Scientific, Pratteln, Switzerland | AC41348 |
| Calcium chloride dihydrate | 0.09 g/L | 0.17 g/L | Merck, Schaffhausen, Switzerland | 102382 |
| Resazurin sodium salt | 1 g/L | 1 g/L | Sigma-Aldrich Chemie GmbH, Buchs, Switzerland | 199303 |
| L-Cysteine hydrochloride monohydrate | 1 g/L | 1 g/L | Acros Organics | 111785000 |

**Table S2:** Supplemented substrates

|  | Product | Supplier | Article No. | Batch No. |
| --- | --- | --- | --- | --- |
| AG | FiberAid larch wood arabino-galactan | Lonza | 175649 | FA16174 |
| PE | Pectin from citrus peel | Sigma-Aldrich Chemie GmbH, Buchs, Switzerland | P9135 | SLBQ6929V |
| XY | Xylan from oat spelt | Chemie Brunschwig AG (Angene), Basel, Switzerland | AGN2018-23001 | G01-23001-1 |
| SS | Starch from potato | Sigma-Aldrich Chemie GmbH, Buchs, Switzerland | S2004 | 0000070988 |
| RD | NUTRIOSE® FB06 | Roquette, Lestrem, France |  | E527N |
| MU | Mucin from porcine stomach (Type II) | Sigma-Aldrich Chemie GmbH, Buchs, Switzerland | M2378 | SLBX8709 |
| YE | Yeast extract for biotechnology Fermtech® | Sigma-Aldrich Chemie GmbH, Buchs, Switzerland | 1.11926 | VM848026823 |
| SU | di-Sodium succinate | VWR International AG, Dietikon, Switzerland | 8.18601 | S7277901841 |
| LT | DL-Lactic acid (90%) | Sigma-Aldrich Chemie GmbH, Buchs, Switzerland | 69785 | BCBPS043V |

## Supplementary Figures

**Figure S1:**
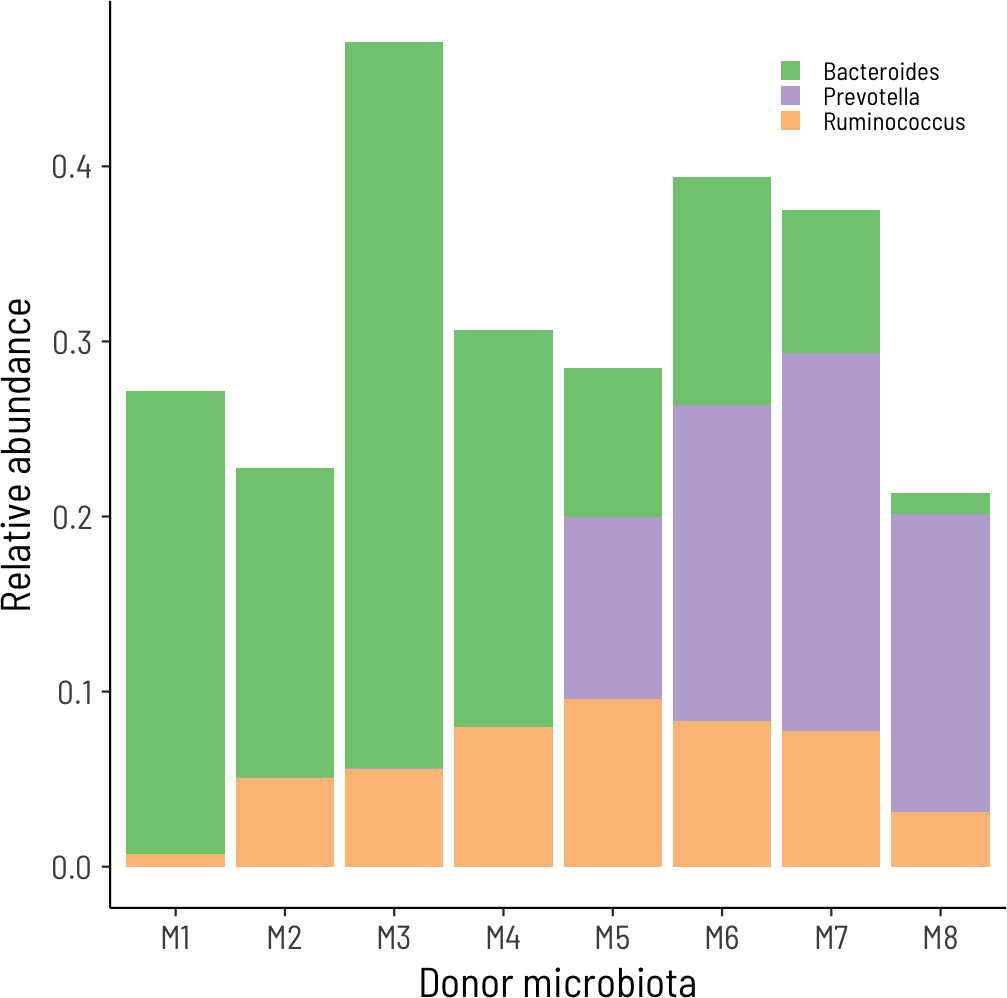
The eight microbiomes span the typical enterotype diversity. For each microbiome, we computed the relative abundance on the genus level. Bacteroides contains both the genera *Bacteroides* and *Phocaeicola*, and Ruminococcus contains all genera *Ruminococcus_A*, …, *Ruminococcus_F*.

**Figure S2:**
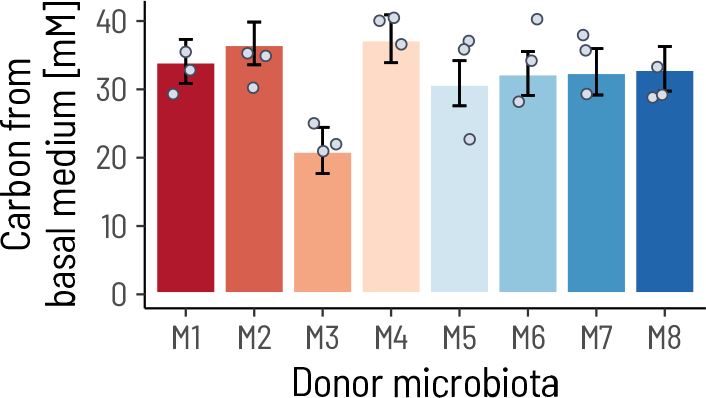
The eight donor microbiota do not differ strongly in their estimated capacity to metabolite the basal medium. The bars show the estimated values, *ξ_d_*, for each donor microbiota and the error bars show the 95% highest probability density interval. The points show the measured total carbon in the measured metabolites. Note, that the estimated values of *ξ_d_* are informed by all of the enrichments and that the model estimated a global measurement error with mean *σ* = 4.75.

**Figure S3:**
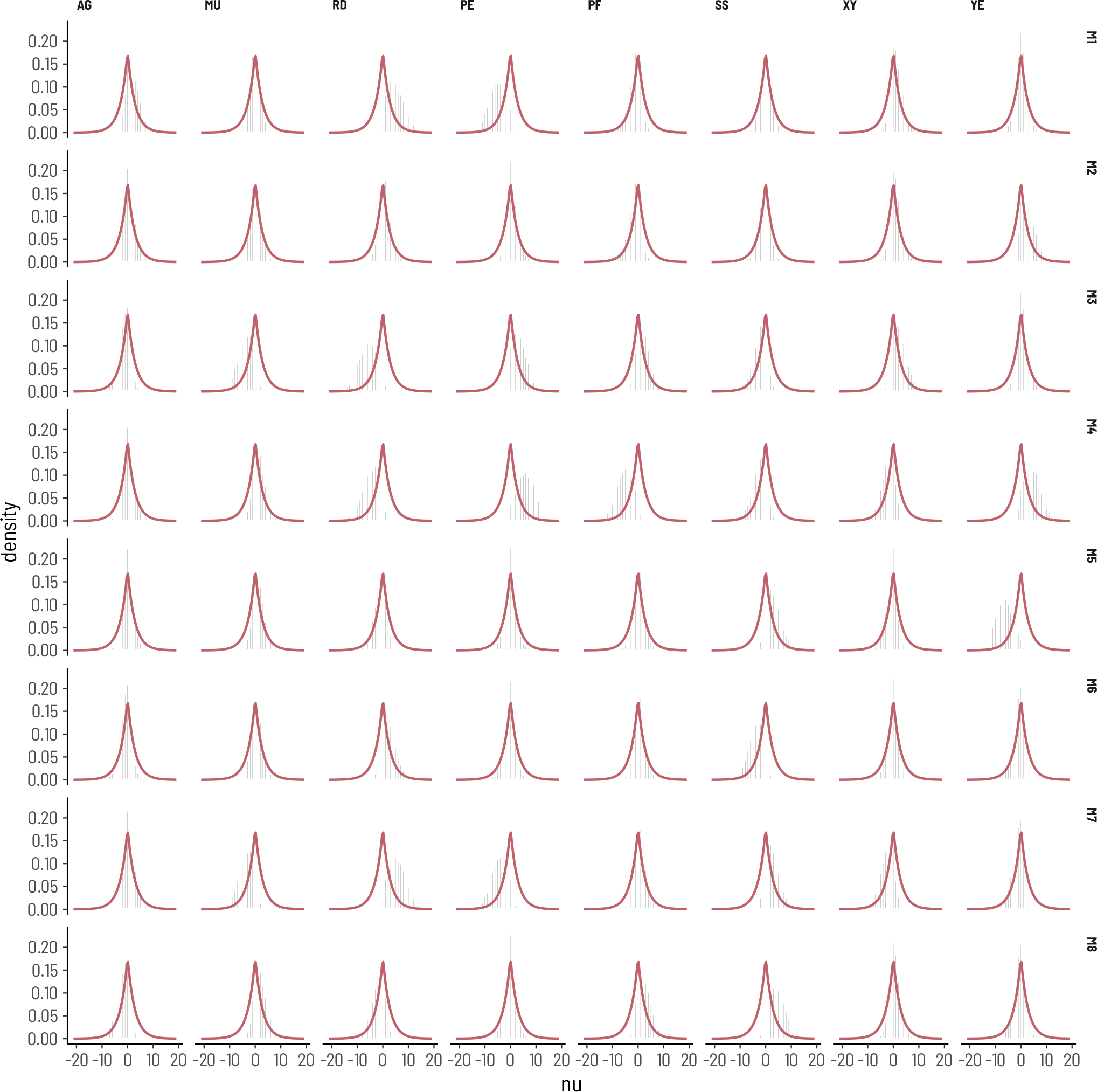
Bayesian posterior density estimates for the parameter *ψ_d,s_*. The bars show the density histograms over 4000 posterior samples. The red lines indicate the Laplace prior using the mean posterior estimate of the scale parameter of the Laplace distribution, *σ_ψ_* = 2.70.

**Figure S4:**
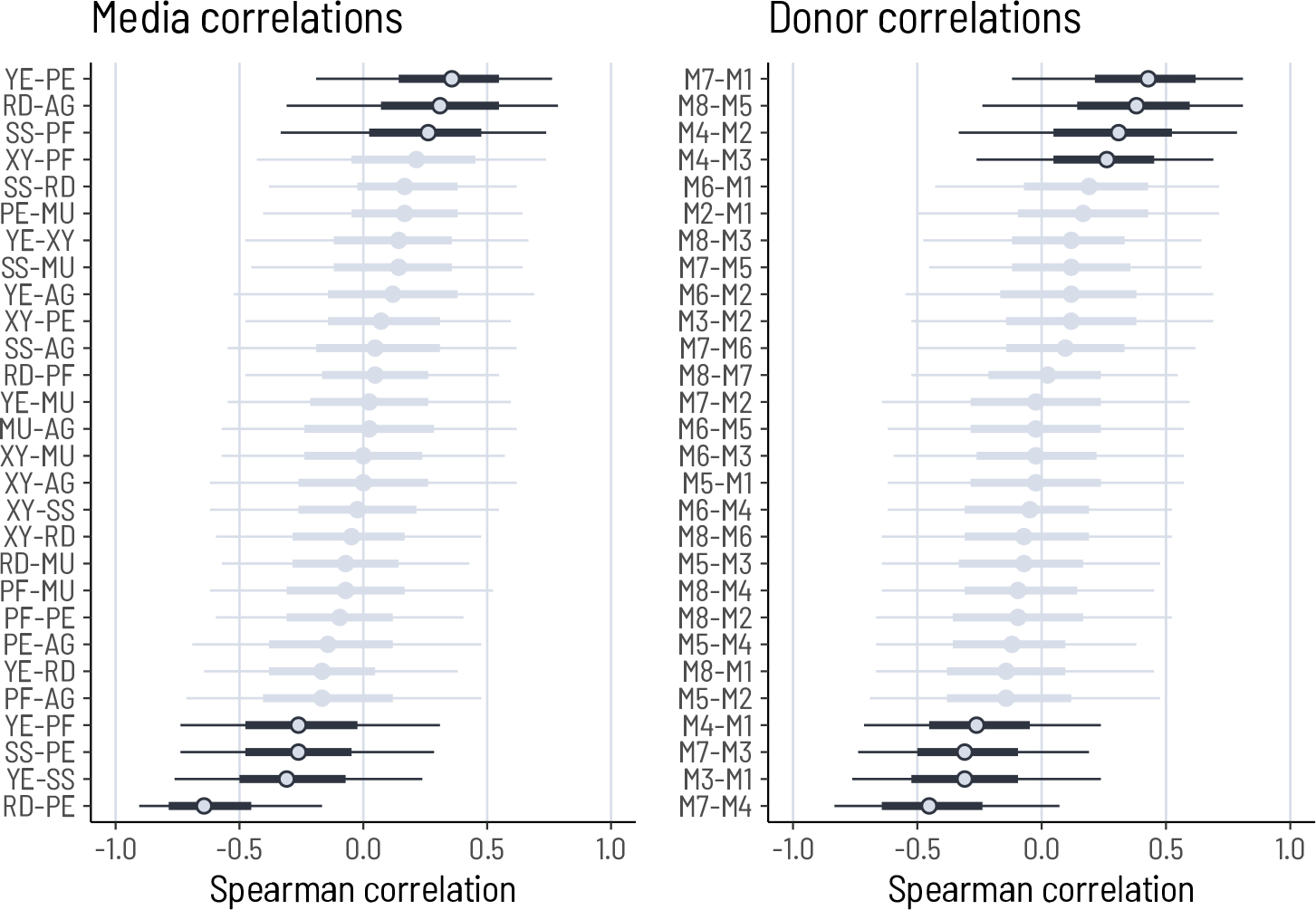
Correlations between donors and metabolites with respect to the *ψ_d,s_*. The circles show the mean of the Spearman correlation coefficients computed for each draw from the posterior. The thick and thin lines show the 50% and 90% inter-quantile ranges (IQR), respectively. Pairs with a 50% IQR that do not overlap with zero are shown in dark.

**Figure S5:**
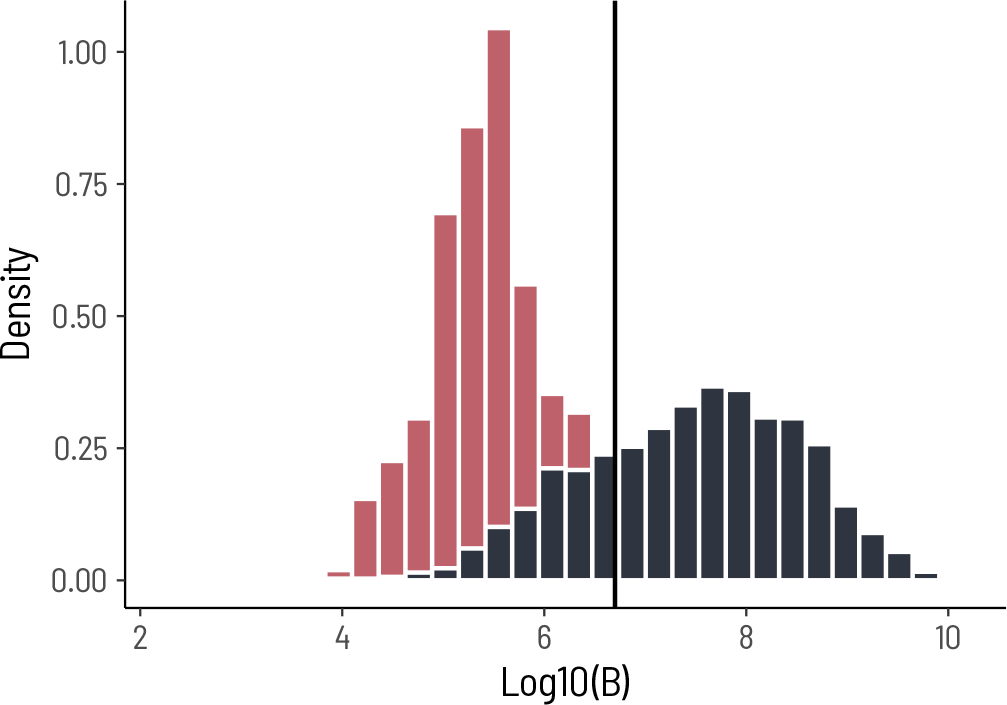
The distribution of absolute growth across all genera and primary substrates is bimodal. The two modes are roughly separated at *B_T_* = 5 *×* 10^6^ cells*/*mL. Red show computed values of *B* that arise solely from the regularization (pseudo count). Thus, only values *B > B_T_* are considered true signals.

**Figure S6:**
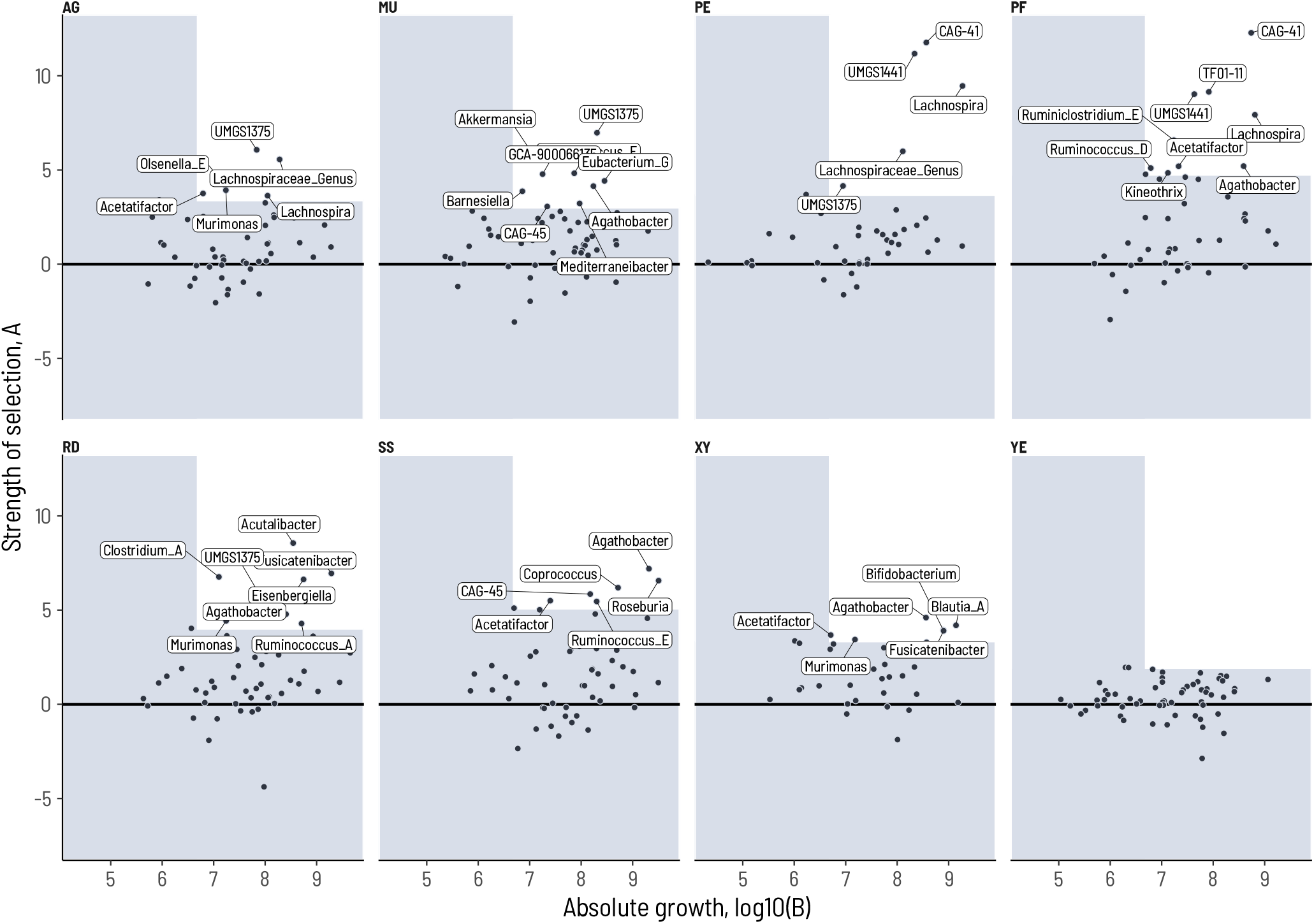
Differentially enriched taxa on primary substrates. For each substrate, we partition the genera into unspecific and specific growth (above/below *B_T_* = 5 *×* 10^6^ cells*/*mL). We then determine the maximum value of *A* for the genera with unspecific growth, *A_T_*. The white area shows those genera with *A > A_T_* and *B > B_T_*.

**Figure S7:**
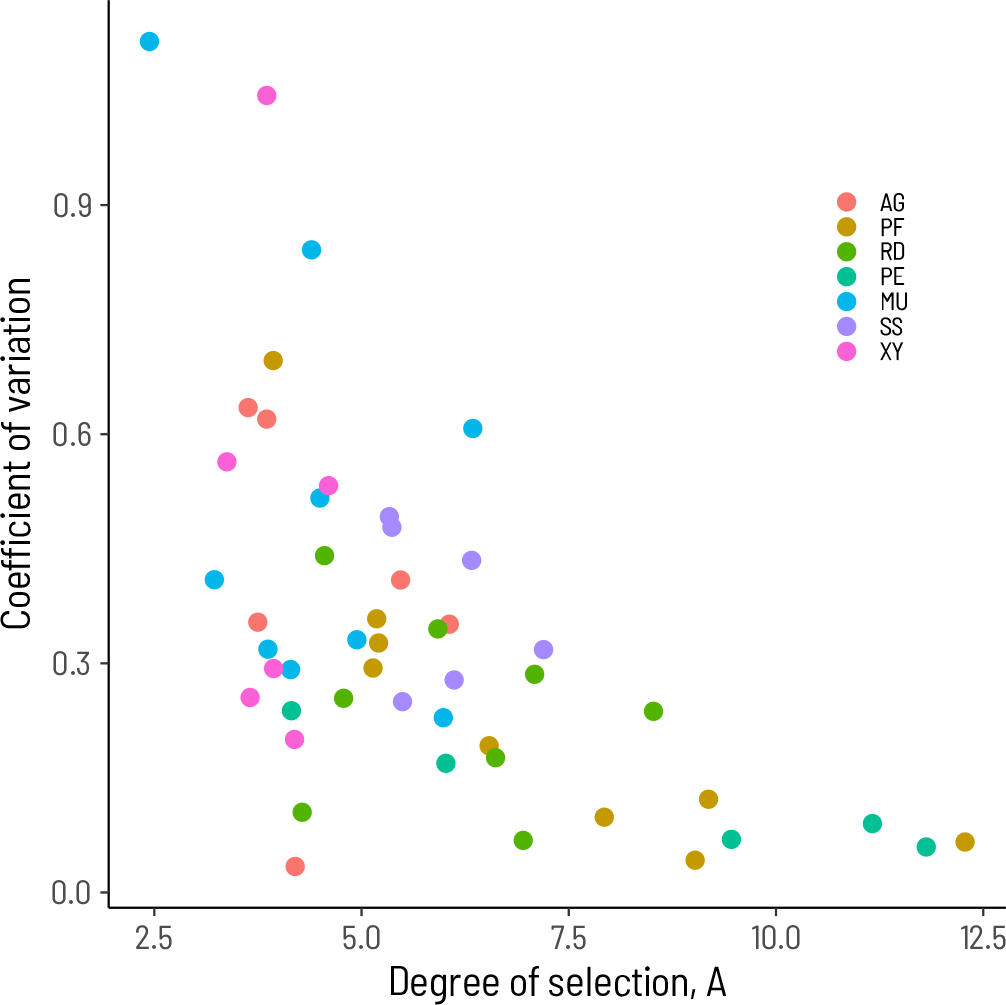
Degree of selection versus coefficient of variation across fecal donors.

**Figure S8:**
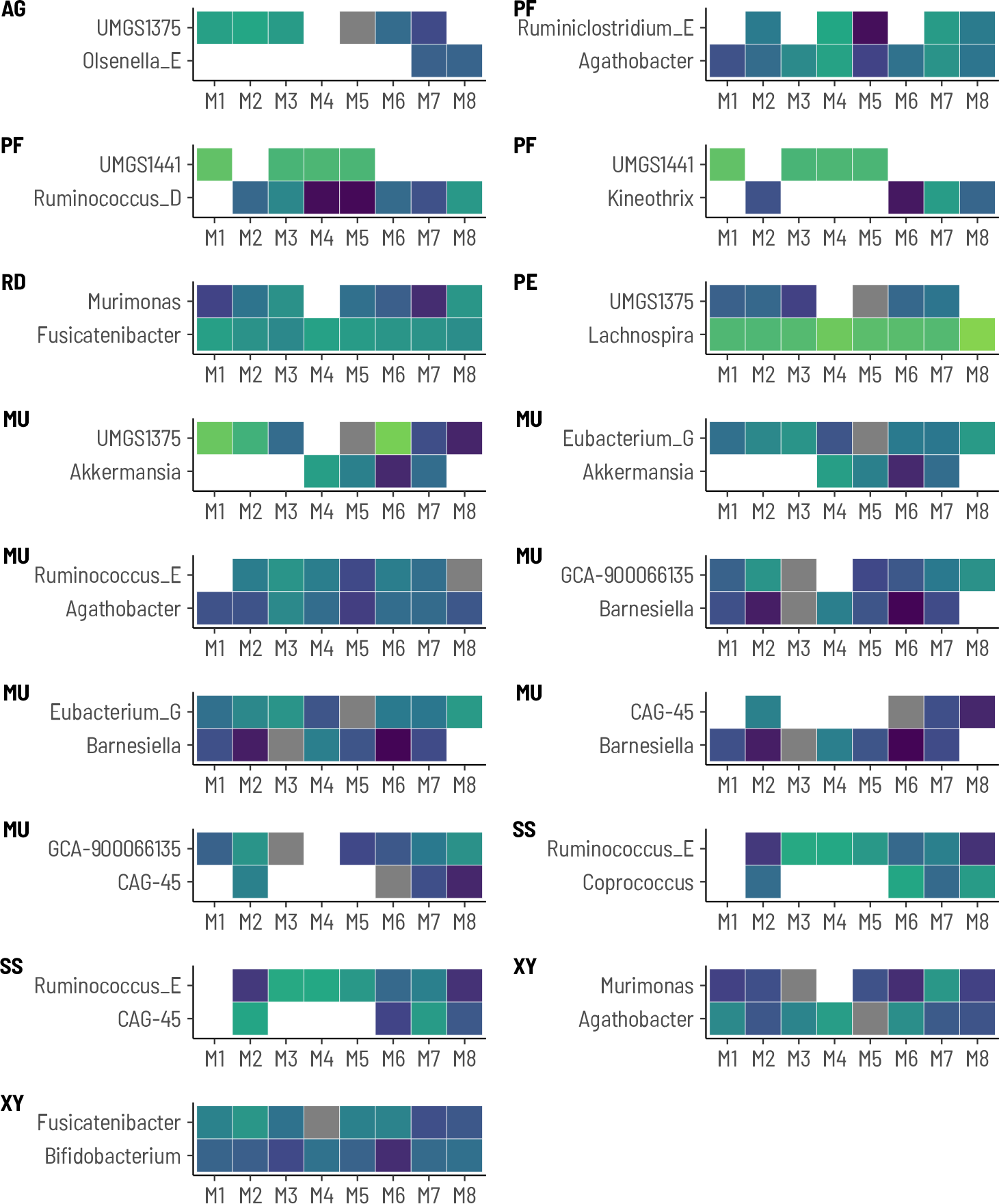
Visualization of the significant correlations between the characteristic taxa. The color scale is the same as in Figure 2.

**Figure S9:**
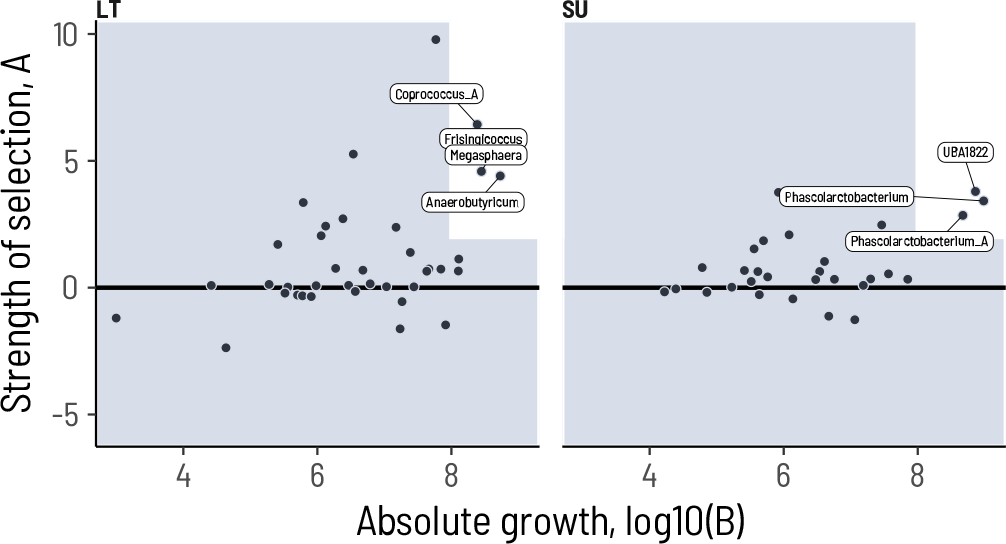
Characteristic taxa for the intermediate fermentation products lactate (LT) and succinate (SU).

**Figure S10:**
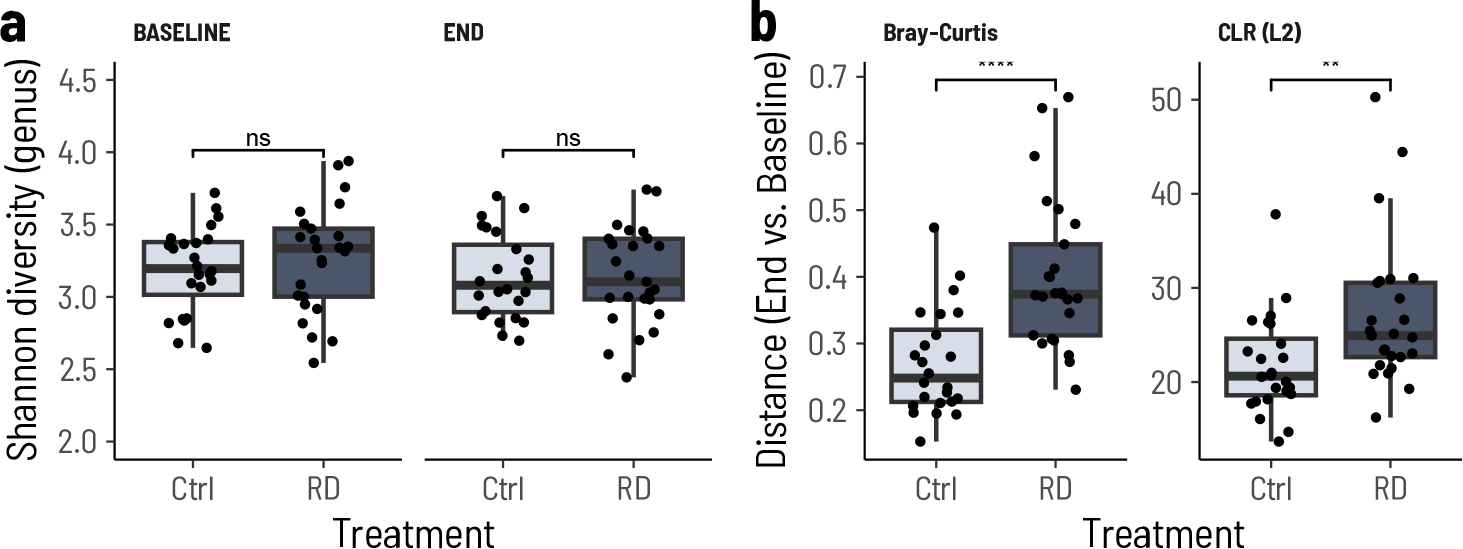
Fecal microbiota changes after dietary supplementation with a resistant dextrin. **a.** Fecal microbiota Shannon diversity of participants in the nutritional intervention study. The Shannon diversity was computed from the genus-level relative abundances. Baseline: *χ*^2^ = 0.577, *p* = 0.447; End: *χ*^2^ = 0.0064, *p* = 0.936. **b.** Change in microbiota composition from Baseline to End as a function of treatment. Each panel shows a different distance metric. Bray-Curtis is half the L_1_ distance on relative abundances. Center log-ratio (CLR) is the L_2_ (Euclidian) distance on CLR transformed relative abundances. Groups were compared with a Kruskal-Wallis test. Bray-Curtis, *χ*^2^ = 17.3, *p* = 0.0000318; CLR, *χ*^2^ = 8.64, *p* = 0.00328. RD: resistant dextrin (NUTRIOSE®), Ctrl: placebo control (maltodextrin).

**Figure S11:**
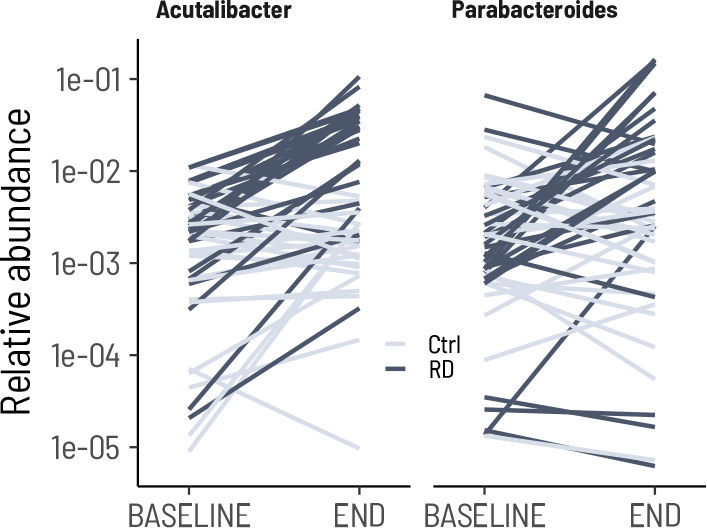
Relative abundance changes for *Acutalibacter and Parabacteroides* between the baseline and end point fecal samples in the nutritional intervention study. Each line represents an individual study participant. Dark lines are the RD group (NUTRIOSE®), light lines are the Control group. The relative abundance is computed including a pseudo count of 1.

## Supplementary Text: Testing for bias from the inoculum

Despite designing our culture conditions to be as broadly suitable for intestinal bacteria as possible, we nevertheless expected a residual bias from laboratory conditions. Furthermore, the dilution at inoculation of 3 *×* 10*^−^*^9^ v*/*v is a strong bottleneck that disfavors rare taxa. To ascertain the degree of enrichment bias, we first looked at how growth in the basal medium without added substrate (mM2) changed microbiota composition.

All eight microbiomes reached cell concentrations over 2 *×* 10^9^ cells*/*mL after 48 h of growth in mM2 and fell into two groups, with 6/8 reaching around 3*×*10^9^ cells*/*mL and 2/8 reaching lower cell densities of 2*×*10^9^ cells*/*mL (Supplementary Figure ST1). Importantly, the density in the fecal sample—and hence the inoculum density—did not correlate with estimated cell densities after 48 h growth in mM2. We thus concluded that differences in cell densities in the feces were not the drivers of the differences at the end of the enrichments.

All eight microbiomes evenly lost diversity after 48 h of culture in the basal medium without supplemented carbon source (Supplementary Figure ST2). We found a strong correlation between the estimated diversity at inoculum and the diversity after enrichment in mM2 (*r* = 0.770, *p* = 0.025). This suggests that the selectivity of the culture conditions *per se* was not systematically biased across microbiomes. To quantify how much of this reduction in diversity resulted from losing taxa while diluting the inoculum, we probabilistically estimated the overall diversity of the microbiota after dilution. Diversity was retained despite the strong dilution (Supplementary Figure ST2a). Hence, dilution at inoculum is an unlikely explanation of diversity loss. Furthermore, there was no significant correlation between the inoculum diversity and the diversity following enrichment for any of the growth conditions (Supplementary Table ST1). We thus subsumed that the specific effect of the growth substrate was sufficiently strong to identify specific taxa.

The composition of the enrichment cultures retained a strong donor-specific signal but showed a consistent shift across all substrates in a principal component analysis (Supplementary Figure ST3). This donor-specific signal was even more apparent when visualizing the change in composition from the fecal sample to the enrichment culture (Main Figure 1g). To correct for this donor bias, we computed the change in composition relative to the composition at the end of the enrichment in the minimal culture medium mM2 (Supplementary Figure ST4). The residual donor signal was much smaller in magnitude compared to the change from the fecal inoculum, allowing us subsequently to identify bacteria-niche associations across donors.

**Figure ST1:**
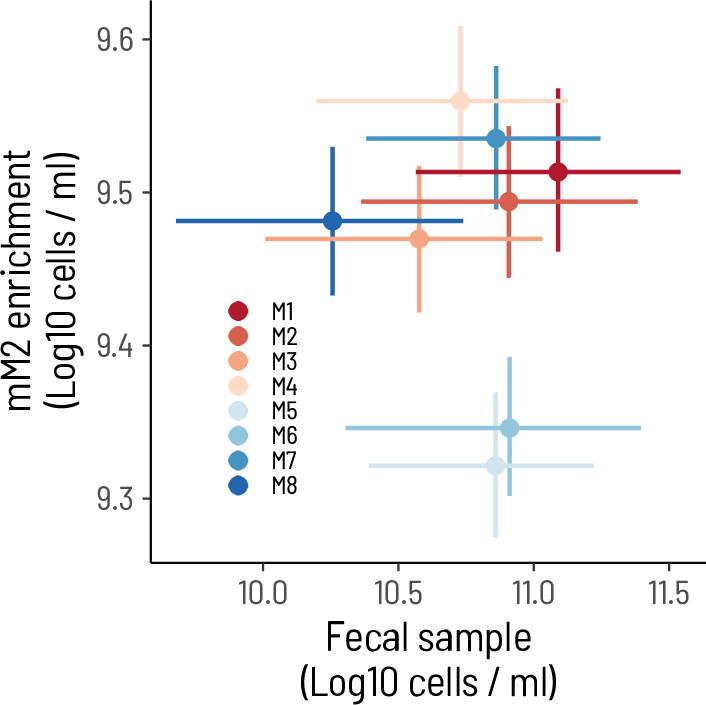
Cell densities in mM2 enrichments do not depend on the density in the fecal sample. Densities in fecal samples were estimated using the MPN method. Densities in mM2 enrichments were estimated from total extracted DNA concentrations calibrated with qPCR quantification with universal 16S primers.

**Figure ST2:**
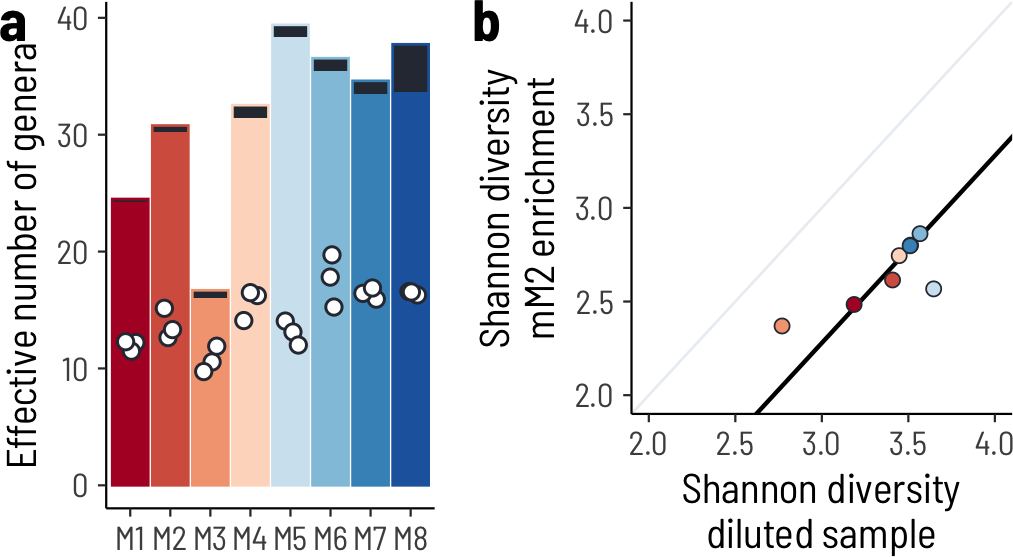
Diversity is evenly decreased in enrichments. **a.** The compositional diversity is only weakly affected by diluting the inoculum. Each colored bar shows the estimated diversity in the diluted fecal samples computed from the estimated cell densities (MPN) and the composition (16S amplicon sequencing). The black bars stacked on top are the additional diversity observed in the respective undiluted fecal sample. The white circles show the diversity that remains after 48 h of culturing in the basal medium without supplemented carbon source. **b.** The diversity is evenly reduced across donor microbiomes after 48 h of culturing in the basal medium without supplemented carbon source. The black line shows a regression excluding microbiomes M4 and M5 and fixing the slope to 1, resulting in an estimate of the intercept of *−*0.72 (CI = [*−*0.76*, −*0.68]).

**Table ST1:** Correlation between inoculum diversity and diversity after enrichment.

| Substrate | Correlation | *p* | Low | High |
| --- | --- | --- | --- | --- |
| AG | 0.40 | 0.32 | -0.42 | 0.86 |
| MU | 0.48 | 0.23 | -0.34 | 0.88 |
| PE | -0.22 | 0.60 | -0.80 | 0.57 |
| PF | 0.49 | 0.22 | -0.33 | 0.89 |
| RD | -0.26 | 0.53 | -0.82 | 0.54 |
| SS | -0.07 | 0.88 | -0.74 | 0.67 |
| XY | -0.08 | 0.86 | -0.74 | 0.66 |
| YE | 0.28 | 0.51 | -0.53 | 0.82 |
| mM2 | 0.77 | 0.02 | 0.15 | 0.96 |

**Figure ST3:**
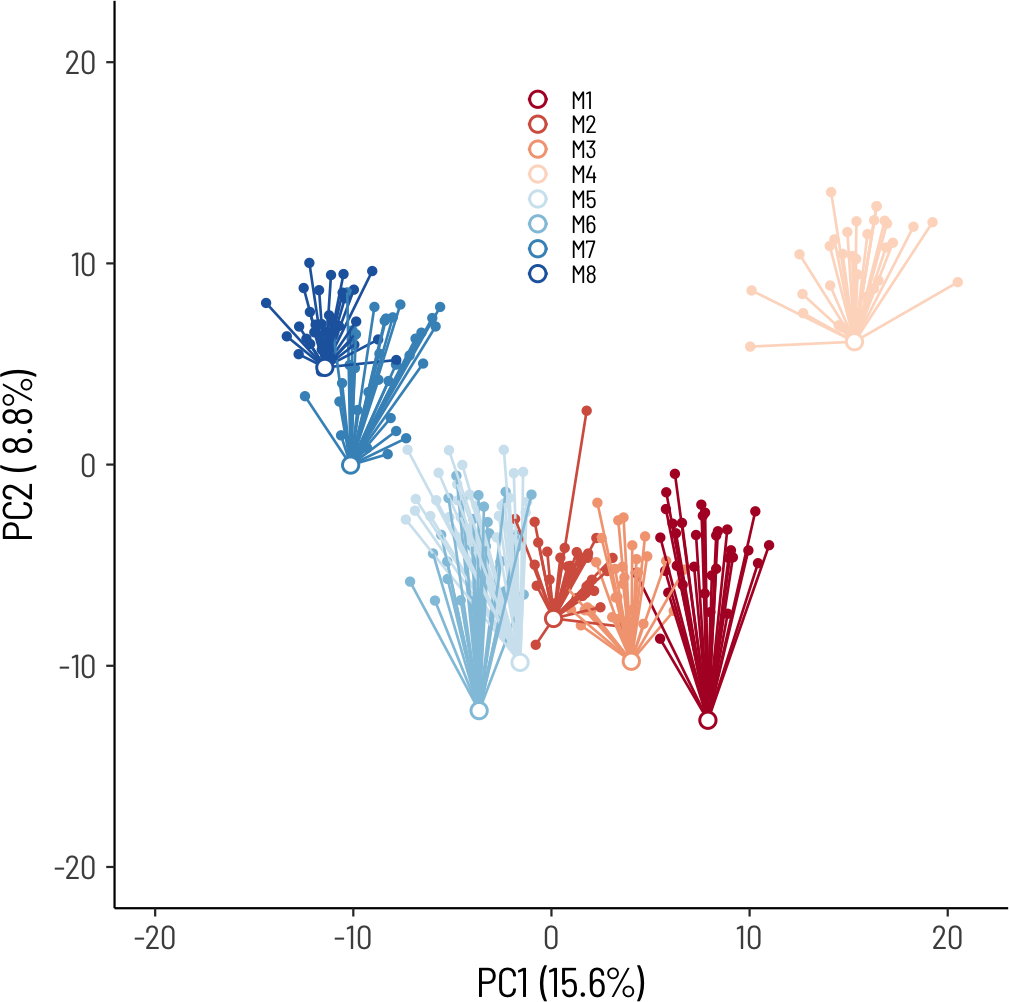
The composition of enrichment cultures retains a signature of the fecal sample. Each point corresponds to a specific enrichment culture. The open white circle shows the composition in the fecal sample. The principle components (PC1, PC2) were computed from the center-log-transformed relative abundances. A pseudo count of 1 was added for each genus.

**Figure ST4:**
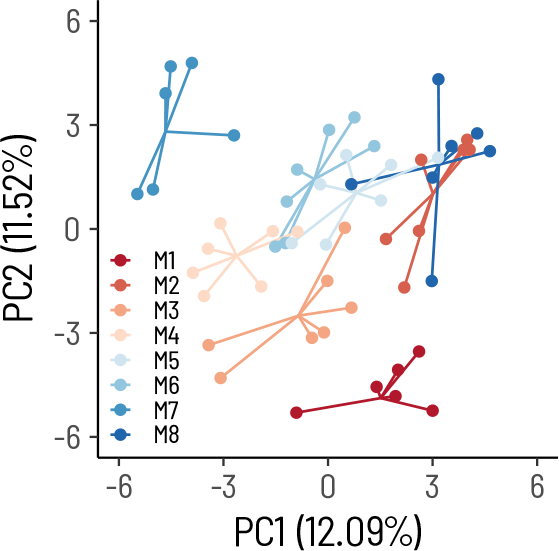
Composition at the end of enrichments after computing the correction relative to mM2. Each point corresponds to a specific enrichment culture.

